# The significance of microRNA genes expression in ovarian cancer cell lines resistant to cytotoxic drugs

**DOI:** 10.1101/859264

**Authors:** Dominika Kazmierczak, Karol Jopek, Karolina Sterzynska, Marcin Rucinski, Radoslaw Januchowski

## Abstract

Ovarian cancer is the most fatal type of gynecological cancer. The main reason for high mortality is the development of drug resistance. It can be related to increased expression of drug transporters and increased expression of extracellular matrix (ECM) proteins. miRNA is a short noncoding RNA that regulates expression of about 60% of genes in the human genome and plays a crucial role in developing cancer, including resistance to chemotherapy.

Our foremost aim was to exhibit alterations in the miRNA expression levels in the cisplatin (CIS), paclitaxel (PAC), doxorubicin (DOX), and topotecan (TOP) - resistant variants of W1 sensitive ovarian cancer cell line using miRNA microarray. The second goal was to identify miRNAs responsible for the regulation of drug-resistant genes.

According to our observation, alterations in the expression of 40 miRNA genes. The level of expression of 21 genes was upregulated in at least one drug-resistant cell line. The expression of 19 genes was downregulated in at least one drug-resistant cell line. We identified target genes for 22 miRNAs. Target analysis showed that miRNA regulates key genes responsible for drug resistance. Among others, we observed regulation of the ABCB1 (MDR1) gene in the W1PR1 cell line by miR-363 and regulation of the COL3A1 gene in the W1TR cell line by miR-29a.

Hence, on the basis of our findings, results indicate that genes responsible for drug resistance development in ovarian cancer can be regulated by miRNA.

## INTRODUCTION

Ovarian cancer poses a growing threat to women’s life and health and occupies the first place in mortality among gynaecological cancers [1]. Most cases of ovarian cancer are diagnosed at a very advanced stage (III or IV according to FIGO) [2,3]. Despite the dynamic development of gynaecologic oncology, the main method of ovarian cancer treatment is the surgical treatment followed by chemotherapy [4]. The first line treatment is based on combination platinum derivatives (carboplatin or cisplatin) and taxanes (paclitaxel - PAC)[2]. The basis for choosing the second line chemotherapy regimen is sensitivity to treatment with platinum derivatives, which determines the prognosis [2]. Low chemotherapy effectiveness result from primary or developed during treatment drug resistance [5]. In over 80% of patients, it comes to recurrence cancer [2]. There are two main types of mechanism of drug resistance: cellular specific and tissue specific [6]. A cellular mechanism is based on cytostatics active removal from cancer cells. Proteins from ABC family contribute to this phenomenon [7]. Cellular mechanisms also include: repairing damaged DNA, developing point mutations in the genes that encode proteins that bind cytostatic drugs and increasing the activity of anti-apoptotic or pro-survival pathways as well as disrupting apoptotic signalling pathways [4]. On the other hand, the tissue specific mechanisms are associated with: tumour vascularization, cell density in the tumour and expression of extracellular matrix proteins (ECM) [8]. All of them changes drug distribution in tumour tissue decreasing their availability to tumour cells. Additionally, ECM proteins can induce cell adhesion mediated drug resistance (CAM-DR) leading to increased resistance to apoptosis [9]. Although many genes associated with the development of resistance to chemotherapy are known, the mechanisms of their regulation are still poorly understood. One of the ways to regulate gene expression is regulation at mRNA level by small noncoding RNA particles – designated as a micro RNA (miRNA). Short non-coding RNAs were discovered and described for the first time in 1993 by Victor Ambros from Harvard University [10]. miRNAs are short (19-29 nucleotides), single-stranded non-coding RNA molecules, having a phosphate residue (5 ‘end) and a hydroxyl group (3’ end) [11]. They play an important regulatory role in the expression of genes in both animals and plants [11]. In the case of miRNA, the regulation of gene expression takes place at the post-transcriptional level and involves interaction with mRNA [12]. The seed sequence (6-8 nucleotides) located in the 5’ region of the miRNA plays a key role in this process. miRNA seed recognizes the sequence of the target mRNA complementary within the 3’UTR (untranslated region). Depending on the degree of complementarity of the mRNA to the mature miRNA, either the transcript degradation (complete or almost complete complementarity) occurs or the translation is inhibited (incomplete complementarity) [11]. Effect of miRNA expression can be pleiotropic. Modulation of one miRNA expression can affect many different genes responsible for the various mechanisms of cancer [13]. The sequences encoding miRNAs in the human genome can be located both in introns of genes and in non-endogenous areas. Some of the miRNA encoding genes may also be located within exons of other genes. miRNAs are classified in groups called of miRNA families and most of them is transcribed by RNA polymerase II [14]. Family members are characterized by: common origin, evolutionary conservation in sequence, similar mature miRNA/miRNA* (shared functional characteristics or biological function) and conserved mature miRNA-seed-target relationships [15,16]. Very interesting is that miRNA genes in the same miRNA family are non-randomly localized around genes connected with cancer, infectious, immune system, sensory system, neurodegenerative diseases, and development [15]. A set of two or more [17] miRNAs which is transcribed from physically adjacent miRNA genes is called a cluster of miRNA. miRNAs that create the same cluster are transcribed in the same orientation without separation by any transcription unit. Clusters of miRNAs take part in controlling various important cellular processes such as tumour formation, development of lungs, heart and immune system [17]. Changes in miRNA expression have been reported in many diseases including different cancers. A miRNA profiling study showed that the pattern of miRNA expression in cancer tissues compared to normal tissues was very different [18]. miRNA can perform suppressor or oncogenic functions in cancers. As a tumour suppressor, it may inhibit cancers’ growth by negative oncogenes’ regulation — generally, suppressor miRNA expression is decreased in cancer cell [19]. On the contrary, the oncogene miRNAs expression (called “oncomirs”) is increased in tumours. Oncomirs promote tumours development and progression by the mechanism of down-regulating a tumour suppressors or other genes involved in for example cell proliferation or cell differentiation in cancer [19]. In ovarian cancer expression of different suppressors: miR-26b [20], miR-29b [21], miR-30a-5p [22], miR-93-5p [23], miR-106b-5p [24], miR-125b [25]; and oncogenes: miR-183 [26], miR-376a [27], miR-383-5p [28], miR-551b-3p [29], miR-572 [30] was described. Ovarian cancer tissue studies revealed the negative correlation between members of the miR-23 family. High expression level of miR-23a and low expression level of miR-23b were connected with the stage and degree of cancer, the differentiation level and lymph nodes metastases [31]. Much attention is currently being paid to miRNA in the context of drug resistance in ovarian cancer [12,18,19]. Changes in miRNA expression are associated with the development of resistance to chemotherapy in patients [32] as well as in vitro in the sensitive-resistant cell line model [32]. In PAC-resistant patients, downregulation of miR-663 and miR-622 were correlated with better prognosis [32]. Overexpression of miR-647 in the PAC-sensitive patients was correlated with good prognosis and that suggested suppressor function of miR-647 [32]. Changes in miRNA expression were also observed in cell line study. Downregulation of miR-31 expression correlated with taxane resistance in ovarian cancer cell lines [33]. In contrast upregulation of miR-98-5p was observed in cisplatin (CIS) -resistant cell lines [34]. Changes in miRNAs expression were also noted in another cancers. miR-195 expression was downregulated in temozolomid-resistant glioma cells [35] and miR-203 was downregulated in prostate cancer cells resistant to doxorubicin (DOX) [36]. The use of miRNA microarrays to analyse changes in miRNA gene expression is an effective molecular tool for the discovering new miRNA genes involved in drug resistance processes. The present study shows alterations in the miRNA expression levels in the CIS (W1CR), PAC (W1PR1 and W1PR2), DOX (W1DR), and topotecan (TOP) (W1TR) - resistant variants of W1 sensitive ovarian cancer cell line. We also identified a set of target genes that can be responsible for drug resistance in these cell lines.

## MATERIALS AND METHODS

### Reagents

Cisplatin, Doxorubicyn, Topotekan and Paclitaxel were obtained from Sigma (St. Louis, MO, USA). RPMI-1640 medium, fetal bovine serum, penicillin, streptomycin, amphotericin B (25 µg/ml) and L-glutamine were also purchased from Sigma. QIazol Lysys Reagent, miRNeasy Mini Kit and RNeasy MinElute Cleanup Kit were obtained from Qiagen (Hilden, Germany). GeneChip™ miRNA 3.1 Array Strip, FlashTag™ Biotin HSR RNA Labeling Kits, GeneAtlas™ Hybridization, Wash, and Stain Kit for miRNA Arrays were obtained from Affymetrix (Santa Clara, CA, USA).

### Cell lines and cell culture

The human primary ovarian cancer cell line W1 was established from tumour tissue of untreated 54-year old Caucasian female patient diagnosed for serous ovarian adenocarcinoma (G3, FIGO IIIc). Cells grow as a monolayer, present epithelial morphology and adherent growth model. Sublines resistant to cisplatin (W1CR), doxorubicin (W1DR), topotecan (W1TR) and paclitaxel (W1PR1 and W1PR2) were obtained by exposure of the W1 cell line to stepwise increasing drug concentrations. Final concentration of each drug was twofold greater than the concentration in the plasma 2 hours after intravenous administration. The cells were 8-, 10-, 20-, 641- and 967-folds resistant to their selective drugs, respectively, as determined by Cell Proliferation Kit I (MTT) [37]. All cell lines were maintained as monolayer in complete medium [RPMI-1640 medium supplemented with 10% (v/v) fetal bovine serum, 2 pMl-glutamine, penicillin (100 U/ml), streptomycin (100 U/ml) and amphotericin B (25 µg/ml)] at 37°C in a 5% CO2 atmosphere.

### miRNA isolation

miRNA was isolated using a reagents kit from Qiagen according to the manufacturer’s protocol. The RNA was quantified using spectrophotometry by measuring the absorbance values at 260 nm and 280 nm, and the 260/280 nm ratio was used to estimate the level of protein contamination. The 260/280 nm ratios of the samples ranged from 1.8 to 2.0.

### Microarray preparation, hybridization and scanning

MiRNA expression profiling was performed using microarray approach with Applied BiosystemsTM miRNA 3.1 Array Strip (ThermoFisher Scientific, Waltham, MA, USA). Detailed technical procedure was described earlier [38,39]. Each microarray was designed in accordance with the miRBase Release 17 database, including complementary probes for:

1733 human mature miRNA, 2216 human snoRNA, CDBox RNA, H/ACA Box RNA, and scaRNA, 1658 human pre-miRNA. The full procedure for preparing miRNA for hybridization was performed using the FlashTagTM Biotin HSR RNA Labeling Kit (ThermoFisher Scientific, Waltham, MA, USA). 150 ng of previously isolated miRNA was subjected to the poly(A) tailing and biotin ligation procedure, according to the manufacturer’s protocol. Biotin-labeled miRNA were hybridized to Applied BiosystemsTM miRNA 3.1 Array Strip (20 h, 48 °C). Afterwards the microarrays were rinsed and stained according to the technical protocol using the Affymetrix GeneAtlas Fluidics Station (Affymetrix, Santa Clara, CA, USA). The array strips were scanned using an Imaging Station of GeneAtlas System (Thermo Fisher Scientific, MA, USA).

### Microarray analysis and gene screening

The initial analysis of the scanned array strips was performed with Affymetrix GeneAtlas Operating Software (Affymetrix, Santa Clara, CA, USA). The quality of the gene expression data was verified using quality control criteria set by the software. The obtained CEL files from the scanned microarrays were imported for further data analysis using BioConductor libraries from the “R” statistical programming language. The Robust Multiarray Average (RMA) normalization algorithm (implemented in the Affy library) was used to normalize, correct the background and calculate the expression values of all the tested miRNAs [40]. The biological annotation was obtained from the pd.mirna 3.1 library, where annotated data frame object was merged with normalized data set, leading to a complete miRNA data table. Differential expression and statistical evaluation were determined using a linear model for microarray data implemented in the “limma” library [41]. The selection criteria for significantly changed gene expression were based on the difference between folds greater than absolute 5 and p-value after false discovery rate (FDR) correction <0.05. The result of such selection was presented as scatter plots diagram showing the total number of miRNAs, up and down regulated (Fig. 1). Differentially expressed miRNA were also visualized as heat map and tables. Raw data files were deposited in the Gene Expression Omnibus (GEO) repository at the National Center for Biotechnology Information (http://www.ncbi.nlm.nih.gov/geo/) under the GEO accession number GEO: GSE134061.

**Fig. 1.**
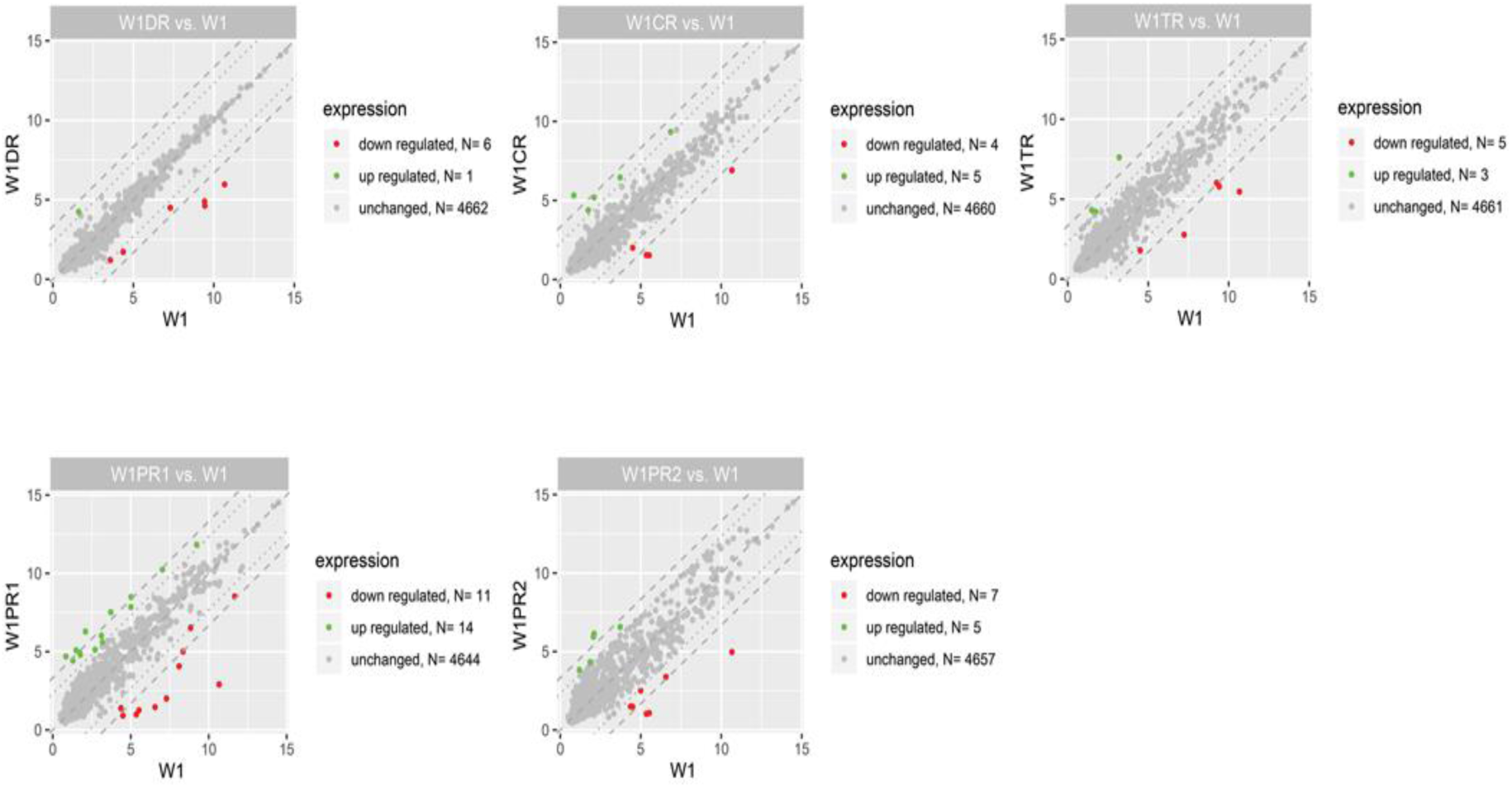
Scater plots displaying the miRNA with expression levels that were up-regulated (green dots) or down-regulated (red dots) by fivefold and more in drug resistant cells in relation to the insensitive W1 cells. Grey dots indicated genes bellow cut-off criteria.

### miRNA-Target Gene Prediction

SpidermiR package was applied to identify potential target genes for differently expressed miRNA. Differentially expressed miRNAs were used as a query for searching target genes in the following databases: for predicted targets—DIANA, Miranda, PicTar, TargetScan, and for experimentally validated targets—miRTAR, miRwalk [42]. To determine the actual expression value of target genes, mRNA transcriptomic data from our published experiment were used [43-45]. Obtained fold change values for mRNA were assigned to target genes data table. For further analysis, we have selected only those target genes for which fold change was inversely correlated with the fold change value of appropriate miRNA (cut-off criteria: fold ± 5, adjusted p-value (adj.p.val.) < 0.05). From the whole set of miRNA-targets pairs, we selcect only those that were involved in drug resistance, extracellular matrix and cancer stem cell biology using a following key words in GO terms: „collagen-containing extracellular matrix”, „extracellular matrix”, „extracellular space”, „response to drug” and „stem cell”. Interactions between miRNA and target genes, in the form selected GO terms, were visualized using Cytoscape 3.7.1 [46].

## RESULTS

### Gene chip quality assessment

In the present study, we used standard factors such as signal to noise ratio internal hybridization, controls spike-in-controls to preliminarily determine the quality of analysed samples. Controls sipke-in-controls: spike_in-control-2, spike_in-control-23, spike_in-control-29, spike_in-control-31, spike_in-control-36. Oligos 2, 23 and 29 are RNA, confirm the poly(A) tailing and ligation. Oligo 31 - poly(A) RNA - confirms ligation. Oligo 36 is poly(day) DNA and confirms ligation and the lack of RNases in the RNA sample.

### Gene expression evaluation and gene expression lists

We evaluated changes in transcription level of miRNA genes. Analysis of the gene expression in five drug resistant ovarian cancer cell lines provided a new information about the significance of changes in miRNA genes expression in drug resistance development in ovarian cancer. Table I summarise the changes in the expression of miRNA genes in drug resistant sublines with respect to drug sensitive W1 cell line. Statistically significant level changed in the drug-resistant cells relative to that in their drug-sensitive counterparts by higher than 5-fold, and less than 0.2-fold (up-/down-regulation of more than/less than 5 and −5, respectively) was evaluated. Genes with expression levels of between 5- and 0.2-fold those of the controls were considered ‘not significant (NS)’ when the gene lists were constructed.

**Table I:** List of the miRNA and the fold changes in the expression of miRNA in each tested line.

| Gene name | miRBase<br>accession number | Fold change |  |  |  |  |
| --- | --- | --- | --- | --- | --- | --- |
|  |  | W1PR1 | W1PR2 | W1CR | W1TR | W1DR |
| hsa-let-7c | MIMAT0000064 | **5,40** | N.S | N.S | N.S | N.S |
| hsa-miR-100 | MIMAT0000098 | **-5,01** | N.S | N.S | N.S | N.S |
| hsa-miR-105 | MIMAT0000102 | **-19,22** | **-21,80** | **-15,92** | N.S | N.S |
| hsa-miR-10a | MIMAT0000253 | **-34,42** | **-9,07** | N.S | N.S | N.S |
| hsa-miR-1271 | MIMAT0005796 | **9,08** | N.S | **6,24** | **5,54** | N.S |
| hsa-miR-1285 | MIMAT0005876 | N.S | **5,54** | N.S | N.S | N.S |
| hsa-miR-143 | MIMAT0000437 | **8,81** | N.S | N.S | N.S | N.S |
| hsa-miR-145 | MIMAT0000437 | **12,07** | N.S | N.S | **7,05** | N.S |
| hsa-miR-146b-5p | MIMAT0002809 | **-39,13** | N.S | N.S | N.S | -6,93 |
| hsa-miR-152 | MIMAT0000438 | **18,19** | **16,52** | **8,53** | N.S | N.S |
| hsa-miR-181a | MIMAT0000256 | N.S | N.S | N.S | **-12,28** | N.S |
| hsa-miR-181b | MIMAT0000257 | N.S | N.S | N.S | **-9,39** | N.S |
| hsa-miR-18b | MIMAT0001412 | **-10,22** | N.S | N.S | N.S | N.S |
| hsa-miR-193a-5p | MIMAT0004614 | **5,32** | N.S | N.S | N.S | N.S |
| hsa-miR-193b | MIMAT0002819 | **7,18** | N.S | N.S | N.S | N.S |
| hsa-miR-195 | MIMAT0000461 | **13,94** | N.S | **6,67** | N.S | N.S |
| hsa-miR-199a-3p | MIMAT0000462 | N.S | N.S | N.S | N.S | **-23,09** |
| hsa-miR-199b-3p | MIMAT0000464 | N.S | N.S | N.S | N.S | **-28,18** |
| hsa-miR-203 | MIMAT0000264 | **14,45** | N.S | **22,33** | N.S | N.S |
| hsa-miR-205 | MIMAT0000465 | N.S | N.S | N.S | N.S | **6,23** |
| hsa-miR-20b | MIMAT0000466 | **-8,87** | N.S | N.S | N.S | N.S |
| hsa-miR-214 | MIMAT0000271 | N.S | N.S | N.S | N.S | **-26,05** |
| hsa-miR-23b | MIMAT0000468 | **6,06** | N.S | N.S | N.S | N.S |
| hsa-miR-23c | MIMAT0000469 | **7,47** | N.S | N.S | N.S | N.S |
| hsa-miR-27b | MIMAT0000470 | **9,21** | N.S | N.S | N.S | N.S |
| hsa-miR-29a | MIMAT0000086 | N.S | N.S | N.S | **-22,04** | N.S |
| hsa-miR-30a | MIMAT0000087 | N.S | **14,82** | N.S | N.S | N.S |
| hsa-miR-31 | MIMAT0000089 | **-217,59** | **-51,72** | **-13,51** | **-36,90** | N.S |
| hsa-miR-335 | MIMAT0000765 | N.S | N.S | N.S | **21,31** | N.S |
| hsa-miR-363 | MIMAT0000707 | **-16,37** | N.S | N.S | N.S | N.S |
| hsa-miR-4269 | MIMAT0016897 | N.S | N.S | N.S | N.S | **-5,13** |
| hsa-miR-449c | MIMAT0016897 | **-11,97** | **-8,09** | **-5,61** | **-6,53** | N.S |
| hsa-miR-4721 | MIMAT0019835 | N.S | **7,18** | N.S | N.S | N.S |
| hsa-miR-497 | MIMAT0002820 | **8,22** | N.S | N.S | N.S | N.S |
| hsa-miR-501-5p | MIMAT0002872 | N.S | **-5,60** | N.S | N.S | N.S |
| hsa-miR-572 | MIMAT0003237 | N.S | **6,23** | N.S | N.S | N.S |
| hsa-miR-708 | MIMAT0004926 | N.S | N.S | **5,52** | N.S | N.S |
| hsa-miR-767-5p | MIMAT0003882 | **-20,61** | **-19,99** | **-13,95** | N.S | N.S |
| hsa-miR-875-3p | MIMAT0004923 | **-7,93** | **-7,32** | N.S | N.S | **-6,28** |
| hsa-miR-99a | MIMAT0000097 | **11,02** | N.S | N.S | N.S | N.S |

### miRNA gene expression in drug resistant cell lines

We observed alterations in the expression of 40 miRNA genes (Table I, Fig. 2). The level of expression of 21 genes was upregulated in at least one drug resistant cell line. Expression of 19 genes were downregulated in at least one drug resistant cell line.

**Fig. 2.**
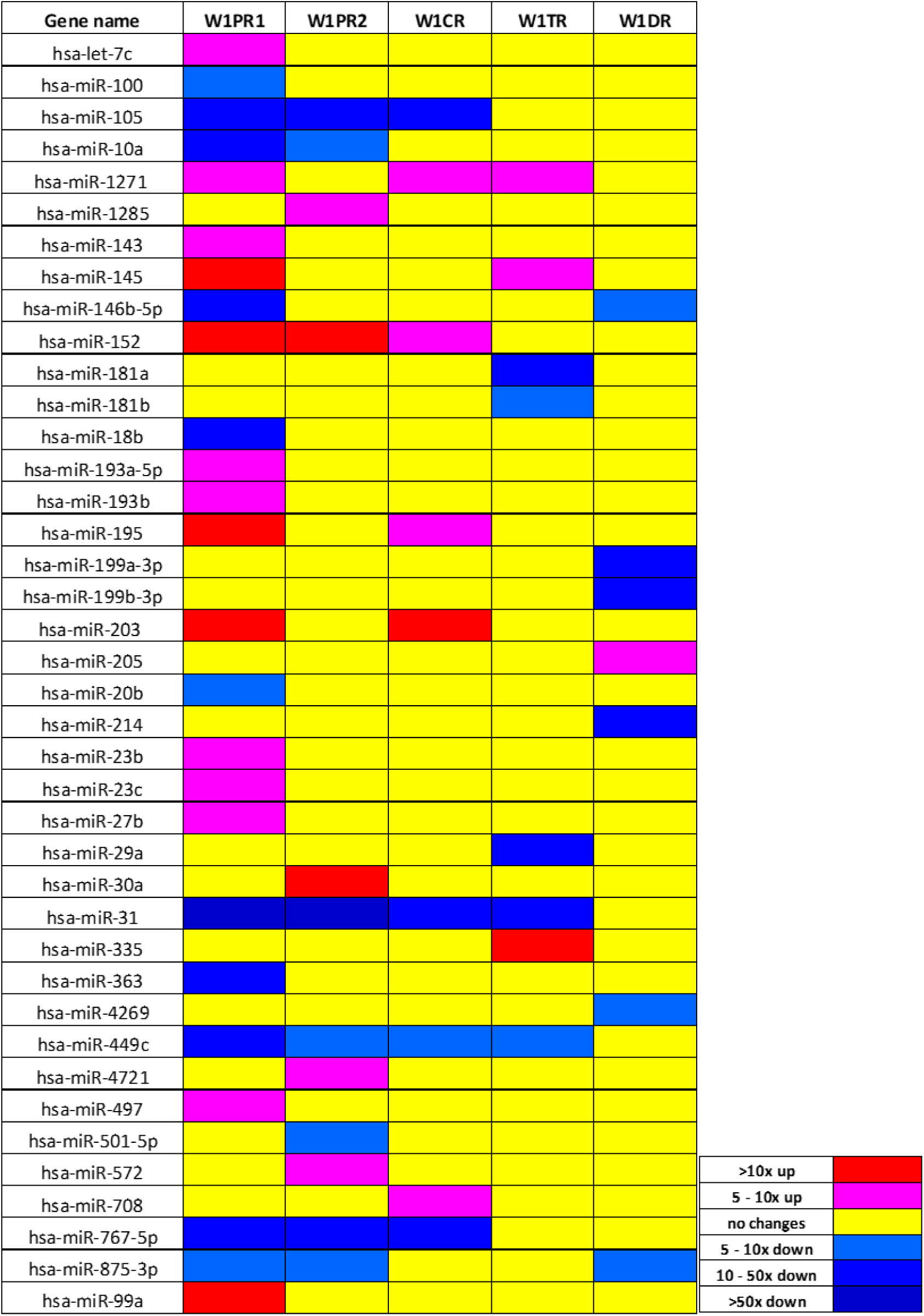
Expression ratios of miRNA genes in drug-resistant sublines.

In relation to the determined cut-off criteria (fold +-5, p<0.05), the most alternations were observed for W1PR1 (PAC resistant cell line) −25 miRNA with altered expression (14 genes upregulated and 11 genes downregulated). Changes in expression of 12 miRNA were observed for W1PR2 cell line with 5 genes upregulated and 7 genes downregulated. Much less changes were observed for other drug resistant cell lines. In CIS resistant cell line, we observed changes in 9 miRNA genes (5 genes upregulated and 4 genes downregulated). In TOP resistant cell line changes in 8 genes expression were observed (3 genes upregulated and 5 genes downregulated) and finally in DOX-resistant cell line we observed changes in expression of 7 genes (1 gene upregulated and 6 genes downregulated).

Based on the following standards, we selected 27 of 40 miRNA genes for further analysis. 1) miRNA genes with changes in expression at least 5-fold (up or down) in at least three drug resistant cell lines; 2) at least 10-fold changes in miRNA gene expression in one drug resistant cell line; 3) 5-fold changes in expression of at least two miRNA genes from the same family.

Expression of miR-31 and miR-449c changed in four drug resistant cell lines (W1PR1, W1PR2, W1CR and W1TR). Expression of miR-105, miR-1271, miR-152, miR-767-5p and miR-875-3p changed in three drug resistant cell lines. Expression of 12 miRNAs was changed at least 10-fold in one tested line, and between them expression of miR-10a, miR-146b-5p, miR-18b, miR-214, miR-29a, miR-363 was downregulated and expression of miR-145, miR-195, miR-203, miR-30a, miR-335 and miR-99a was upregulated.

Expression of miRNA from families 181 (miRNA-181a and miRNA-181b), 193 (miRNA-193a-5p and miRNA-193b), 199 (miRNA-199a-3p and miRNA-199b-3p) and 23 (miRNA-23b and miRNA-23c has also been changed in investigated cell lines. What is important the expression of miRNAs belonging to the same family has been altered in the same way. Especially members of 193 and 23 families were upregulated in W1PR1 cell line and members of 181 and 199 family were downregulated in W1TR and W1DR cell lines, respectively.

High similarity between cell lines resistant to PAC was observed. In both cell lines the same six genes (miR-105, miR-10a, miR-31, miR-449c, miR-767-5p and miR-875-3p) were downregulated and miR-152 was upregulated. Some similarity was also observed between both PAC and CIS resistant cell line – drug used in the first line chemotherapy of ovarian cancer.

Of the 27 genes selected for analysis, nine showed a very significant level of expression change, >20 fold. miR-31 was downregulated in four cell lines (W1PR1, W1PR2, W1CR and W1TR) with very strong downregulation more than 217-fold in W1PR1 cell line and strong downregulation – 52-fold in W1PR2 cell line and 36-fold in W1TR cell line. Among other significantly downregulated genes, we observed downregulation of miRNA-10a (34-fold), miRNA-146b-5p (39-fold) and miRNA-767-5p (20-fold) in W1PR1 cell line. We also observed a significant downregulation of miR-105 in W1PR2 (22-fold) cell line, miRNA-29a (22-fold) in W1TR cell line as well as downregulation of miRNA-199a (23 fold), 199b (28-fold) and miRNA-214 (26 fold) in W1DR cell line. Significant upregulation of miRNA-203 (22-fold) was observed in W1CR cell line and of miRNA-335 (21 fold) in W1TR cell line.

### Analysis of target genes expression

In the second part of our investigation we were interested if described miRNAs are involved in regulation of genes responsible for drug resistance development. By the assumption that increasing miRNA expression leads to decreasing target gene expression and vice versa, for further analysis we selected those target genes in which the fold change was inversely correlated with the fold change of miRNA and changes at least five times up/down respectively with adj. p val<0.05. Targets with expression levels of between 5- and 0.2-fold (up/down regulation between 5 and −5) those of the controls were considered ‘not significant (NS)’ when the target gene lists were constructed. For analysis of target genes expression we used our microarray data that were published previously [43-45]. From target genes analysis we excluded miRNAs expressed only in W1PR2 cell line, because we do not possess mRNA microarray data for this cell line. So, the following miRNAs were excluded from analysis: miR-1285, miR-30a, miR-4721, miR-501-5p, and miR-572. Using our criteria (changes in expression 5 fold up or down) we did not found any targets for the following miRNA: miR-1271, miR-152, miR-203, miR-335 and miR-214.

For further analysis we selected genes involved in drug resistance, extracellular matrix and cancer stem cell biology using a following key words in GO Data Base: response to drug, drug transport, extracellular space, extracellular matrix, collagen containing extracellular matrix, stem cell, because previously we described changes in expression of genes from these groups in investigated drug resistance cell lines (43-45). Using this criteria we did not found targets for the following miRNA: miR-100, miR-10a, miR-143, miR-193a-5p, miR-199a-3p, miR-199b-3p, miR-4269 and miR-99a.

In CIS resistant cell line we identified targets for miR-767-5p, miR-195 and miR-708. Among them miR-195 overexpression correlated with *SLC2A14* (solute carrier) transporter downregulation (Fig.3). In DOX resistant cell line we found targets for miR-146b-5p, miR-205 and miR-875-3p, and among them we could distinguish *Semaphorin 6A* (*SEMA6A*) regulated by miR-875-3p (Fig. 4).

**Fig. 3.**
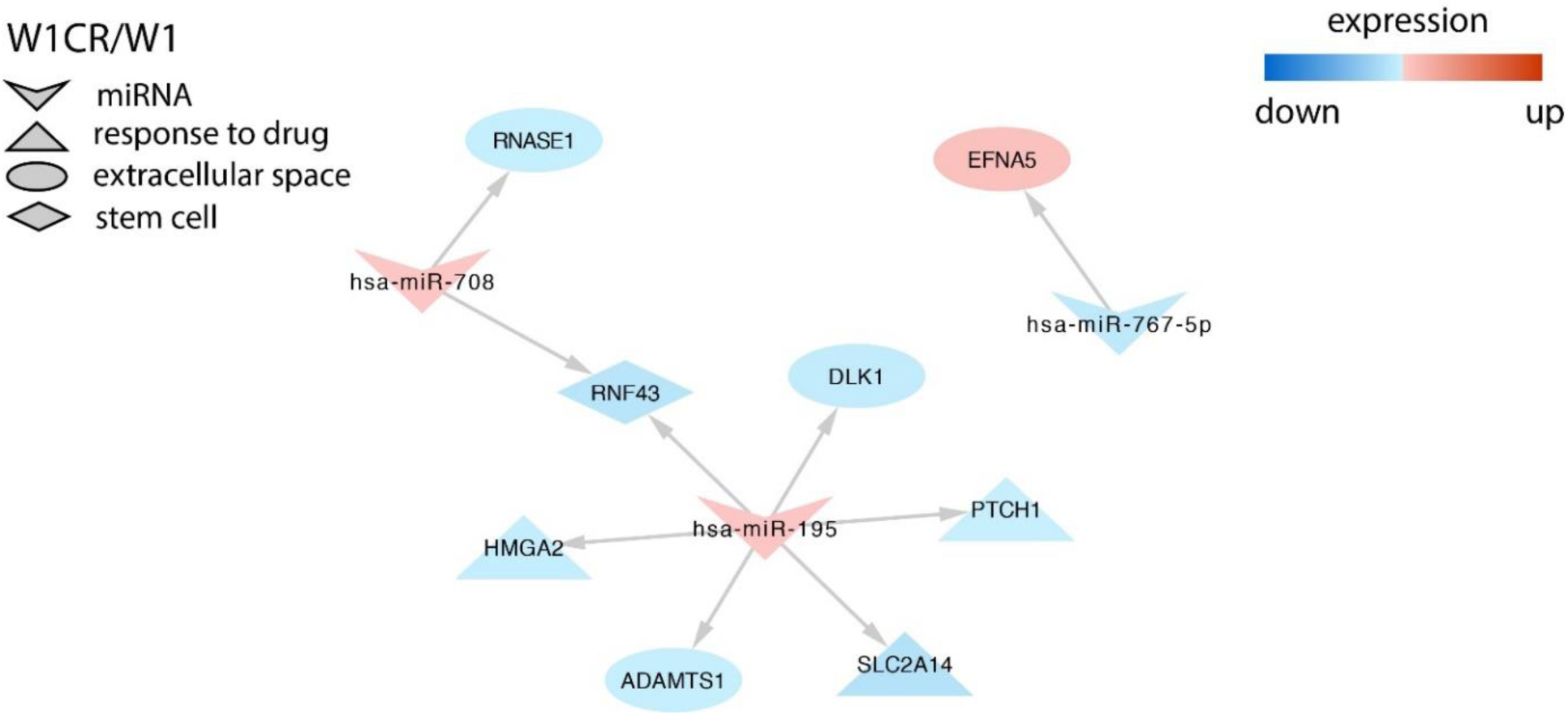
Regulation of selected target genes by miRNAs in the W1CR cell line.

**Fig. 4.**
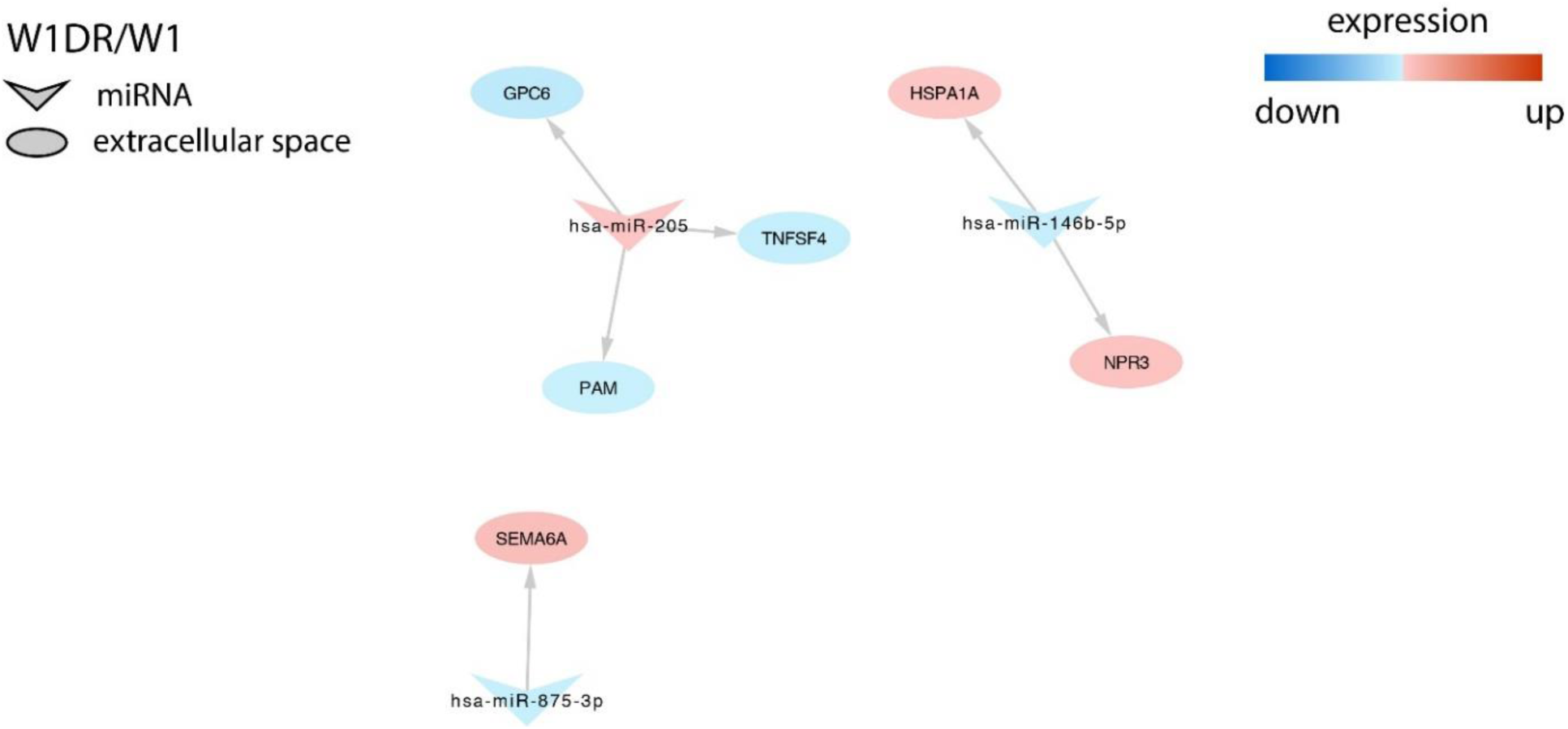
Regulation of selected target genes expression by miRNAs in the W1DR cell line.

Much more targets were identified in PAC and TOP resistant cell lines and among them important genes related to drug resistance or other features of cancer cells. In W1PR1 cell line we observed invers correlation between expression of miR-363 and key drug resistant gene *ABCB1* (*ATP Binding Cassette Subfamily B Member 1*). Expression of important collagens – *COL3A1* (*Collagen Type III Alpha 1 Chain*) and *COL5A2* (*Collagen Type V Alpha 2 Chain*) was correlated with miR-767-5p, overexpression of receptor tyrosine kinases *EPHA7* (*Ephrin Type-A Receptor 7*) was associated with downregulation of miR-18 and miR-20b. miR-20b and miR-146b-5p downregulation also was correlated with upregulation of *MAP3K8* (*Mitogen-Activated Protein Kinase Kinase Kinase 8*). *SEM3A* (*Semaphorin 3A*) and *MYC* (*Myelocytomatosis Viral Oncogene Homolog*) down regulation was associated with overexpression of miR-145. *MYC* expression was also regulated by let-7c. Upregulation of miR-497 and miR-195 was associated with *PCDH9* (*Protocadherin 9*) downregulation (Fig. 5).

**Fig. 5.**
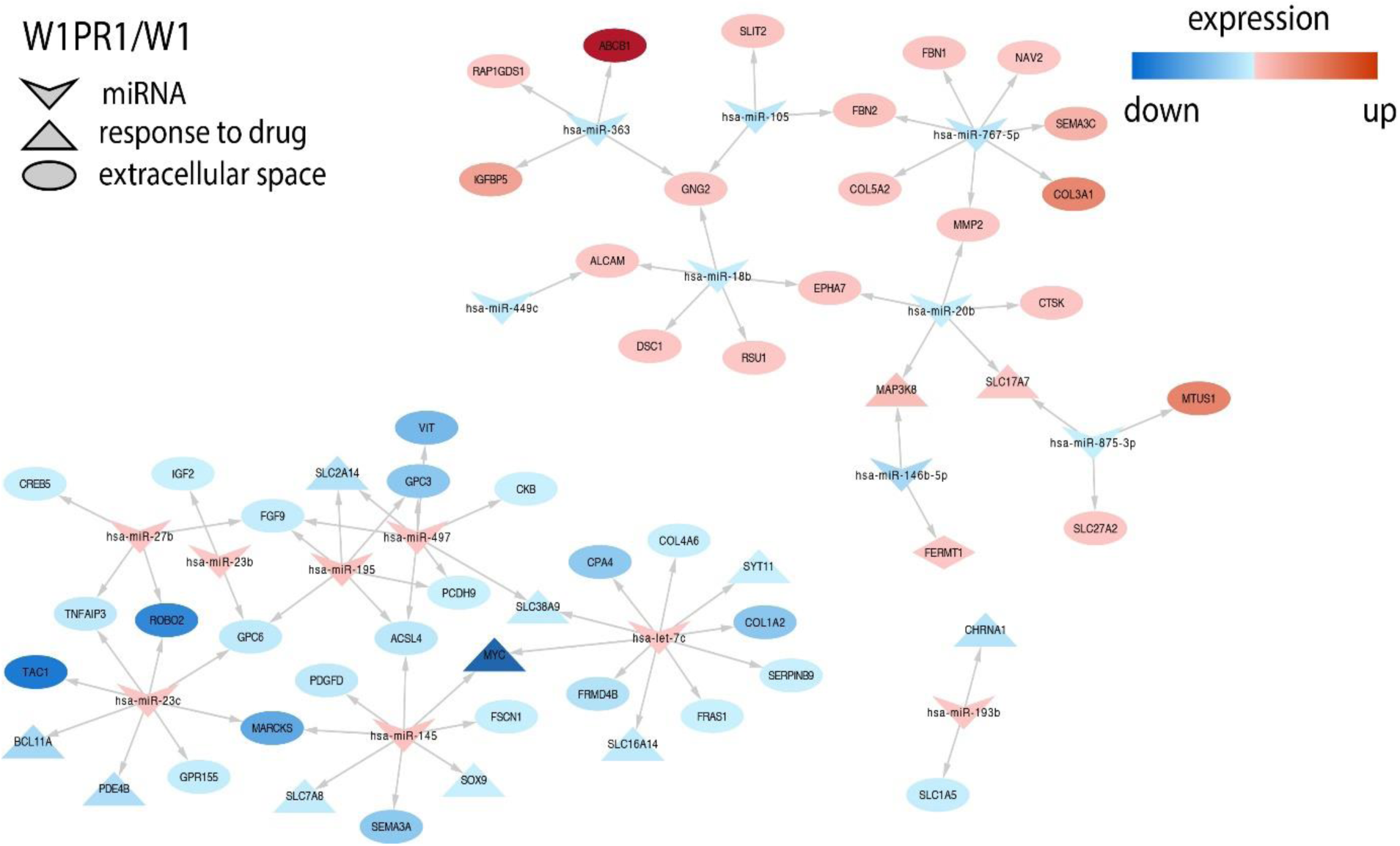
Regulation of selected target genes expression by miRNAs in the W1PR1 cell line.

Changes in different miRNAs were also associated with changes in expression of important drug resistant genes in W1TR cell line. miR-181a and miR-181b downregulation was associated with increased expression of *SPP1/OPN* (*Secreted Phosphoprotein 1/ Osteopontin*), *TGFBI* (*Transforming Growth Factor Beta Induced*), *LOX* (*Lysyl Oxidase*), *EPHA7, DCN* (*Decorin*) and *MAP3K8* (only miR-181b). Expression of *SPP1* was also associated with miR-449c downregulation and miR-449c downregulation correlated with *SEMA3D* (Semaphorin 3D) overexpression. Similarily, in W1PR1 cell line *MYC* downregulation was associated with overexpression of miR-145. Very high upregulation of *COL3A1* was associated with downregulation of miR-29a in this cell line (Fig.6).

**Fig. 6.**
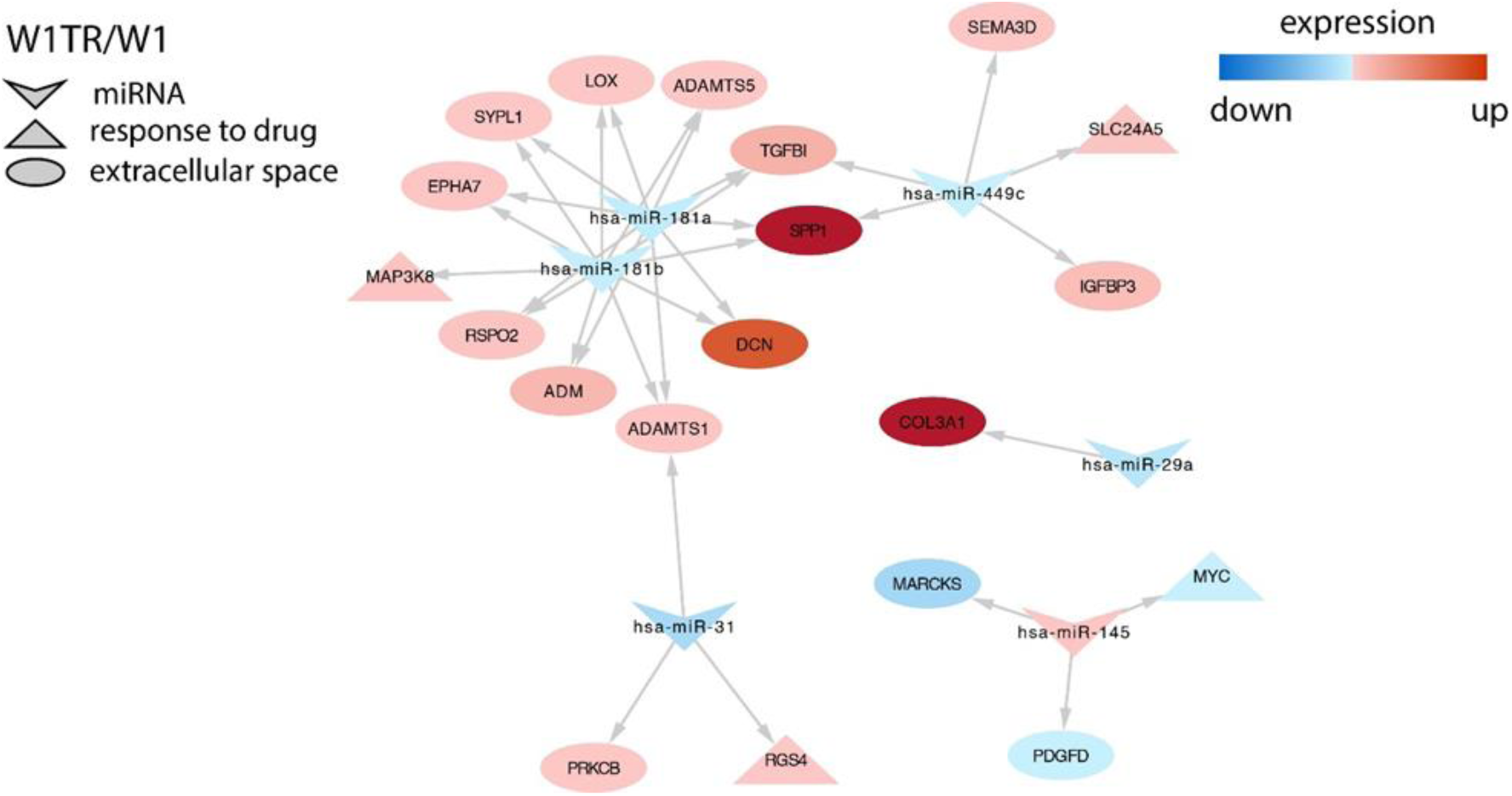
Regulation of selected target genes expression by miRNAs in the W1PR1 cell line.

## Discussion

The present study demonstrates the correlation between resistance to cytotoxic drugs and expression of genes encoding micro RNA. Genes up- or down-regulated less than 5-fold were considered as not significantly altered and not taken into consideration. Between genes that are up- or down-regulated between 5- and 10-fold we consider only those that are changed in at least three drug resistant cell lines or coordinate expression of two genes from the same family is observed. All genes with expression up- or down-regulated more then 10-fold are considered as an important in drug resistance development.

It has been reported that changes in miRNA genes expression can be related to development of chemotherapy resistance in solid tumours as well as in development of resistance of cancer cell lines to cytotoxic drugs in vitro [47]. Thus, in present study we analysed changes in expression of several miRNAs in ovarian cancer cell lines resistant to drugs from the first line (CIS and PAC) as well as from second line of chemotherapy (DOX and TOP). We were especially interested in changes of miRNA genes expression in cancer cells exposed to drugs with different mechanism of action. In contrast to studies in which only one pair of sensitive/resistant cell lines is tested, in this study changes in miRNA expression in five drug resistant cell lines among which PAC-resistant twin cell lines were examined. It is a unique model for such research.

Among the 27 miRNAs analysed in this experiment, we could observe significant downregulation of miR-31 and miR-449c in four out of five drug resistant cell lines. Such a significant decrease in expression in cell lines resistant to CIS, TOP and, which is noteworthy, in both PAC resistant cell lines, may indicate a role in the process of resistance to these drugs. This is supported by other studies in which a downregulation of miRNA-31 induced taxane resistance in ovarian cancer cells. miR-31 was downregulated in a PAC resistant cell line, and reintroduction of miR-31 again sensitized them to PAC both in vitro and in vivo [33]. The miR-31 suppressor effect in ovarian cancer cell line and tissue has been confirmed by Hassan M.H. et al. [48]. They observed a high level of miR-31 in the chemosensitive serous ovarian cancer cell line and decrease in taxane-resistant line [48]. All these results suggest that downregulation of miR-31 expression may be associated with PAC resistance in ovarian cancer. The potential role of reduced miR-31 expression in CIS resistance can also be assumed. This is demonstrated by the results of our study, in which significant downregulation in CIS-resistant cell line was noted. This is in line with the results of other studies that showed decreased miR-31 levels in CIS-resistant tumours and cell line of gallbladder cancer (GBC). In contrast, the ectopic expression of miR-31 in a CIS-resistant cell line causes an increased CIS sensitivity [49]. Because in our study, for the first time, we observed downregulation of miR-31 expression in a TOP-resistant cell line, the potential contribution of miR-31 to the development of TOP resistance requires further investigation.

Unlike miR-31, expression of miR-449c has not been described in the literature in the context of drug resistance in cancer. Since we have demonstrated a downregulation of miR-449c in cell lines resistant to different cytotoxic agents, this may suggest its non-specific role in drug resistance. Previously, we observed ALDH+ stem cell like population in W1PR1, W1TR and W1PR2 cell lines [50,51]. Here, we observed downregulation of miR-31 and miR-449c in all these cell lines, and importantly, the level of these miRNAs downregulation was inversely correlated with the number of ALDH+ cell population. This suggest the role of miR-31 and miR-449c downregulation in development of stem cell population. Lv C. et al. showed that miR-31 plays a critical role in mammary stem cell self-renewal and breast tumorigenesis by regulating Wnt pathway [52].

Increased expression of miR-1271 in our PAC-, CIS- and TOP-resistant cell lines contradicts the results of other investigators. Expression of miR-1271 was downregulated in gastric cancer and cell lines in comparison to healthy tissues and cells and further downregulated in CIS-resistant cell lines [53]. Glioblastoma patients with low level of miR-1271 have shorter overall survival than patients with higher level [54].

We have demonstrated significant downregulation of three subsequent miRNAs: miR-105, miR– 767–5p in CIS and PAC-resistant and miR-875-3p in PAC and TOP-resistant cell lines. Lu G. et al. showed that in NSCLC low miR-105 expression was associated with poor patient survival, suggesting its hypothetical role in chemotherapy resistance [55]. In contrast, in triple negative breast cancer, it was shown that miR-105 was upregulated and correlated with poor patient survival and promoted chemoresistance [56]. However, clinical study does not reflect a simple drug sensitive/resistant cell line model. In contrast, we did not found any data about the role of miR-767-5p and miR-875-3p in drug resistance. Thus, the role of miR– 767–5p, miR-875-3p in drug resistance can be considered possible, but drawing specific conclusions requires more thorough research.

Upregulation of two another miRNAs (miR-152 and miR-195) in cell lines resistant to CIS and PAC could indicate their involvement in the regulation of genes associated with resistance to first line chemotherapy of ovarian cancer. However, in contrast to our study downregulation of miR-152 and miR-195-5p was observed in CIS-resistant ovarian cancer cell lines [57,58]. On the other hand, the upregulation of miR-195 has been observed in PAC-resistant laryngeal cancer cell line [59] and docetaxel resistant head and neck squamous cell carcinoma [60]. Thus, the role of this miRNAs in drug resistance could be cell and drug type dependent.

Based on the second research hypothesis assumed in advance, a group of miRNAs whose expression level was changed at least 10-fold in one drug resistant cell line was selected. In this group, as many as nine different miRNAs (miR–10a, miR–146b–5p, miR-203, miR-30a, miR–363, miR–18b, miR–145, miR-195 and miR–99a) showed changes in expression level in PAC resistant cell lines. However, we could also observe high changes in expression of single miRNAs in CIS- (miR-195, miR-203), TOP- (miR–29a, miR-145, miR-335) and DOX-resistant (miR-146b-5p, miR-214) cell lines. We are aware that the obtained results cannot clearly indicate the role of these molecules in the development of drug resistance. Some of them were described here for the first time (miR-99a, miR-145 and miR-18b). For others, we searched any literature data describing their involvement in the context of drug resistance or, if not found, at least in the pathogenesis of various cancers.

As the result of literature search we found that other research teams described the changes in expression levels of the same miRNAs as ours. In contrast to our result, where decreased level of miR-10a in PAC-resistant cell line was noted, Sun W. et al. observed increased expression level in CIS-resistant A549 lung cancer cell line. Furthermore, miR-10a silencing, increases sensitivity of resistance cell line to CIS [61]. These differences can result from different drugs and different cancer type used in both study. Further studies report that low level of miR–146b– 5p expression correlates with ovarian cancer progression. In advanced cancer stage (III/IV) the level of miR-146b-5p expression was lower compared to earlier stage (I, II) [62]. In cell line study overexpression of miR-146b-5p enhanced ovarian cancer cell sensitivity to PAC and CIS [62]. Thus, miR-146b-5p downregulation seems to play a role in resistance of ovarian cancer to cytotoxic drugs, especially to PAC.

In opposite to our result (upregulation in PAC and CIS resistant cell lines) Cheng R. et al. shown that in CIS-resistant NSCLC tumours miR–203 was downregulated compare with CIS-sensitive tumours [63]. Ectopic expression of miR–203 in NSCLC cell line increases CIS sensitivity and apoptosis [63]. Overexpression of miR-203 also reversed PAC-resistance in colorectal cancer [64]. Regarding the miR-30a, the increase in expression of which we observed PAC-resistant cell line, Sestito et al. miR-30a was downregulated in CIS- and PAC-resistant ovarian cancer cell lines [22] and overexpression of miR-30a caused increased sensitivity of ovarian cancer cells to CIS-induced apoptosis [22].

In contrast to our report, Mohamed Z. et al. [65] observed upregulation of miR-363 in PAC resistance ovarian cancer cell line. Furthermore, overexpression of miR–363 confers PAC resistance and inhibition of miR–363 restores the response to PAC [65]. In contrast downregulation of miR-363 was observed in CIS-resistant hepatocellular carcinoma cell line [66]. Significantly lower expression level of miR-363 was also observed by Cao L. et al. in CIS-resistant ovarian cancer cell line [67]. Thus, the significance of miR-363 in drug resistance can be cell line dependent.

The next three miRNAs described in this report hypothetize their involvement in resistance to DOX, CIS and TOP. In ovarian cancer cell line study downregulation of miR-29a correlated with increased CIS resistance, while overexpression of miR–29a correlated with sensitized ovarian cancer cell line to CIS [68]. Similar results were obtained by Yang L. et al. in glioblastoma stem cells in which overexpression of miR-29a in CD133+ GBM (glioblastoma multiforme) stem cells effectively reversed the resistance to CIS [69]. Downregulation of miR-29a was also observed in methotrexate resistant osteosarcoma cell line [70]. Here, we observe downregulation of miR–29a in TOP-resistant cell line. Thus, it appears that a downregulation of miR-29a could be involved in cytotoxic drug resistance.

In our study miR-335 is significantly upregulated in TOP-resistant cell line. In contrast, downregulation of miR-335 was observed in PAC and CIS resistant A2780 ovarian cancer cell lines [71]. In DOX-resistant cell line we could observe very significant downregulation of miR-214. Downregulation of this miR was also observed in DOX-resistant urothelial bladder cancer tissues and cells lines [72]. Thus, the miR-214 can by a potential marker of DOX resistance. We for the first time observe upregulation of miR-99a, miR-145 and downregulation of miR-18b in PAC-resistant cell line. Upregulation of miR-99a was observed in CIS-resistant gastric cancer cell lines [73] and vincristine-resistant acute lymphoblastic leukemia (ALL) [74]. In contrast to our study, downregulation of miR-145 was observed in PAC-resistant variants of A2780 and SKOV-3 ovarian cancer cell lines [75]. We did not find any information about the significance of miR-18b in drug resistance in general. Thus, its role in drug resistance require further investigation.

It is assumed that miRNAs from the same family can play a very similar role in regulation of genes expression. Therefore, as the last step in our research, we analysed changes in miRNAs expression in the same families. In our study miR-193a-5p miR-193b are upregulated in PAC-resistant cell line that suggest they can regulate expression of gene involved in resistance to this drug. As demonstrated by Khordadmehr et al., the miR-193 family (miR-193a-3p, miR-193a-5p, miR-193b3p, and miR-193b-5p) plays an important role in the pathogenesis of ovarian cancer [76]. A similar result was obtained for miR-23 family members - miR-23b and miR-23c. As there is no literature data describing the contribution of these miRNAs as a cluster to cytostatic resistance, at this stage one can only assume that they may affect the expression of genes associated with PAC resistance. However, the upregulation of miR-23b correlated with better survival of lung cancer patients [77] and on the other hand with DOX-resistance in thyroid cancer cells [78]. In another study downregulation of miR-23b and miR-27b was observed in MDR Ehrlich ascites tumor cells [79]. Thus, the role of these miRs in drug resistance require further investigation. On the other hand, the downregulation of miR-181a and miR-181b in TOP-resistant ovarian cell line, might be an indicator of their involvement in development of TOP resistance. However, their role in TOP-resistance have not been investigated by others. Decreased expression of miR-181a was observed in CIS-resistant cervical cancer cell line [80] and miR-181b in NSCLC cell CIS-resistant cell line [81].

Members of miR-199 family were described in ovarian cancer. Cheng W. et al. showed that miR-199a-3p and miR-199a-5p overexpression significantly decrease chemoresistance of cancer-initiating cells (CICs) to CIS, PAC, and DOX and reduced mRNA expression of the multidrug resistance gene *ABCG2* in vivo [82]. miR-199b-5p silencing is significantly associated with acquired chemoresistance in ovarian cancer cell line and cancer tissue [83]. Here we observed significant downregulation of miR-199a-3p and miR-199b-3p in a DOX-resistant cell line. However, as previously reported, *ABCG2* gene expression has not changed in this cell line [43]. In contrast, this cell line was characterized by overexpression of the *MDR1* gene. Determining the involvement of miR-199 family members in the regulation of drug resistant genes require further investigation.

As presented above, both our results and studies of other researchers indicate some involvement of specific miRNAs in the process of drug resistance. However, the interpretation of miRNA expression results is much more complicated compared to gene encoding proteins. As an example, the expression of *MDR1* gene encoding P-gp protein is always related to drug resistant phenotype. In the case of miRNA genes, this phenomenon more comprehensive. On the one hand, one miRNA gene can regulate the expression of different mRNAs, and from other hand, the same mRNA can be regulated by different miRNAs. This generates a complex network of dependencies. Another aspect that affects the complexity of those relationships is related to the fact that different genes are involved in the drug resistance. From our experience in working on several drug sensitive/resistant pairs of cell lines and based on literature data, we know that some specific genes are involved in resistance to a particular cytostatics, e.g. MDR1 is very often expressed in PAC and DOX resistant cell lines, and *ABCG2* gene, encoding BCRP is very often expressed in TOP-resistant cell lines [45,84]. However, in addition to these canonical genes of drug resistance, there are many other genes whose altered expression is observed in drug resistance, among them e.g. genes encoding extracellular matrix proteins (ECM) that seems to be involved in cell adhesion mediated drug resistance (CAM-DR) [44,85].

Considering the high complexity of the process of acquiring resistance by cancer cells, and summarizing the results obtained in this study, we believe that the contribution of miR-31, miR-146b, miR-195 in PAC-, miR-214, miR-199a-3p and miR-199a-5p in DOX resistance is highly possible. However, the role of the described miRNA genes requires further molecular, cellular and bioinformatics analysis, which we intend to proceed in the next step of our study.

To shed a more light on the role of investigated miRs in drug resistance in the second part of our study we analysed correlation between expression of miRs and their targets. We were mainly interested if genes described by ours previously in the context of drug resistance can be regulated by miRNAs, although some other genes are also discussed.

In our previous microarray analysis [43-45,84,85] we could observe much more genes with changed expression in PAC- and TOP-resistant cell lines than in CIS- and DOX-resistant ones. This seems also to be reflected by number of targets to miRNAs identified in GEO database in these cell lines. Thus, according to number of identified targets we can divide our cell lines into those with low number of identified targets – represented by CIS- and DOX-resistant cell lines and those with high number of identified targets, represented by PAC- and TOP-resistant cell lines.

In W1CR1 cell line the only gene that was described in context of CIS-resistance was *SLC2A14* regulated by miR-195 [45]. The only gene described in context of drug resistance in DOX-resistant cell line was SEMA6A regulated by miR-875-3p. However, in contrast to our results downregulation of *SEMA6A* was observed in DOX-resistant A2780 ovarian cancer cell line [86]. Thus, the role of this miR and its target in drug resistance require further investigation.

As we described previously [37] the most important gene responsible for PAC resistance in W1PR1 cell line was *ABCB1* (*MDR1*) encoding the glycoprotein P (P-gp). Here we observe that upregulation of this gene is related to downregulation of miR-363 in this cell line, that is the first such observation. However, downregulation of miR-363 was associated with *ABCC1* overexpression in oxoplatin hepatocellular carcinoma cell lines [87].

Previously we reported overexpression of several collagen genes in different drug resistant ovarian cancer cell lines [88]. In this experiment, we could observe that increased expression of *COL3A1* and *COL5A2* inversely correlates with miR-767-5p level. Thus, expression of this two genes seems to be regulated by this miRNA. Similarily, we observed downregulation of miR-767-5p in W1PR2 cell line accompanied by very strong *COL3A1* upregulation [88]. This further support the possible role of miR-767-5p in *COL3A1* gene regulation.

Drug resistant cells are characterized by strong signal transduction. Previously we reported increased total pTYR level in drug resistant cell lines in comparison to parental cell lines W1 [89]. Here, we observe that upregulation of receptor tyrosine kinase – *EPHA7* and *MAP3K8* kinase is inversely correlated with miR-18b (*EPHA7*) and miR-18b and miR-20b (*EPHA7* and *MAP3K8*). Regulation of EPHA7 expression by miR-18b was also described by others [90].

It has been reported that signal transduction by EPHA7 receptor may activate ERK signalling pathway that also involves MAP3K8 [91]. Upregulation of EPHA7 was observed in many cancers including hepatocellular carcinoma where it was associated with tumour progression, invasiveness and metastasis [92]. MAP3K8 was described as a mediator of vemurafenib resistance of thyroid cancer stem cells [93].

SEMA3A is a member of semaphorin family of proteins that play a role in many developmental processes but also in cancer progression [94]. It is a tumour suppressor gene and its downregulation correlates with progression of gastric [95] and ovarian cancer [96]. Recently we described *SEMA3A* downregulation in three PAC-resistant ovarian cancer cell lines [97]. Here we showed that expression of *SEMA3A* can be influenced by miR-145 upregulation in PAC resistant cell line. Regulation of *SEMA3A* by miR-145 is not surprising because SEMA3A as a direct target of miR-145 has been described recently [98]. The upregulation of miR-145 was also discovered as connected with strong downregulation of Myc Proto-Oncogene Protein in W1PR1 cell lines. miR-145 has been described as a suppressor miRNA that directly target *MYC* transcript in esophageal squamous cell carcinoma [99] and ovarian cancer [100]. Moreover, as our analysis shows, the *MYC* downregulation can be due to let-7c upregulation in this cell line as well, that was described previously in prostate cancer cells [101].

Another gene, found to be downregulated in PAC resistant cell line was *PCDH9*, the expression on which correlated with miR-195 and miR-497 upregulation. PCDH9 (protocadherin 9) a member of protocadherin protein family [102] is a calium-dependent adhesion protein playing role in neural cell interaction [103]. It is a tumour suppressor gene with decreased expression in ovarian [104] and gastric cancer [105] and its downregulation usually correlates with disease progression and shorter survival. Previously we observed the *PCDH9* downregulation in three PAC resistant cell lines [97] suggesting its significance in resistance to this cytotoxic agent. However, there are no literature data available describing the regulation of *PCDH9* gene by miR-195 or miR-497.

The very important role in TOP-resistant cell line play downregulation of miR-181a and miR-181b, since their downregulation was inversely correlated with upregulation of key genes involved in drug resistance. As in W1PR1 cell line also in W1TR cell line we observed increased total pTYR level [89]. Additionally, increased *EPHA7* and *MAP3K8* level was noted as well. It can be associated with stronger signal transduction, increased pTYR level and eventually drug resistance. As opposed to W1PR1 cell line both genes are regulated here by miR-181a and miR-181b, which is the first such observation described so far.

We could also observe that both - miR-181a and miR-181b - regulate *DCN* and *LOX* genes’ expression. DCN (decorin) is a small leucine reach proteoglycan, a part of ECM that bind to collagens and play an important role in cancer development and metastasis [106]. Regulation of *DCN* by miR-181 has been reported in skin and wound healing [107]. LOX (lysyl oxidase), is a secretory protein involved in cross-linking of collagens and elastin in the ECM that results in increase ECM stabilization [108]. Its expression seems to be an important metastatic factor in breast [109] and ovarian [110] cancer among others. Previously we described increased expression of LOX in PAC and TOP resistant ovarian cancer cell lines and its expression was upregulated in ALDH1A1 positive cells [51]. However, the regulation of *LOX* gene by miR-181a or miR-181b was not described in literature so far.

Two other genes important in drug resistance and cancer progression were also regulated by miR from 181 family as well as miR-449c. TGFBI (transforming growth factor-beta-induced protein), is a collagen I, II and IV binding protein present in ECM. It has been described as the metastasis promoting protein in ovarian cancer [111] and its expression was correlated with shorter survival of patients with serous ovarian cancer [112]. Recently we also described TGFBI as a gene related to TOP resistance in three different ovarian cancer cell lines [43]. It has been proved that expression of TGFBI is regulated by miR-181a in osteoblast during differentiation [113].

SPP1 (secreted phosphoprotein 1), also known as osteopontin (OPN), was originally described in bone tissue [114], however its expression was also described in different cancers including ovarian [115], as a protein involved in tumor progression, metastasis and drug resistance [116]. It has been reported that PAC-, DOX- and CIS-resistance was related to induction of drug transporters protein expression by SPP1 [117] or blocking caspase [118]. Recently we also described expression of SPP1 protein in three TOP resistant cell lines [43]. Similar to our results, the downregulation of miR-181a and upregulation of *OPN* mRNA were observed in CIS-resistant cervical cancer cells [80] and overexpression of miR-181a led to reduced expression of *OPN* and higher sensitivity to CIS. Regulation of *OPN* by miR-181b was described also in eosinophilia in astma [119]. In contrast regulation of *SPP1* by miR-449c has not been described so far.

As we described previously W1TR cell line was characterized by abundant *COL3A1* expression, that was secreted from the cells [88]. *COL3A1* was also described as a protein involved in CIS resistance in ovarian cancer [120]. Here we observed that upregulation of *COL3A1* inversely correlates with downregulation of miR-29a. In similar way upregulation of *COL3A1* was observed in two MTX resistant osteosarcoma cell lines and correlated with miR-29a downregulation. Furthermore, it has been proved that miR-29a regulate *COL3A1* expression [70].

In summary, we identified a set of potential or already described targets for miRs with altered expression in drug resistant cell lines. Some of these targets have previously been described as an important factor in development of drug resistance and/or cancer progression. The way of regulation of some of these target genes by miRs described in our study was previously described. Regulation of other targets require detailed study at molecular level and significance of this regulation need to be confirmed in context of drug resistance.

## References

1. Roett, M.A.; Evans, P. Ovarian cancer: an overview. American family physician 2009, 80, 609–616.

2. Hennessy, B.T.; Coleman, R.L.; Markman, M. Ovarian cancer. Lancet (London, England) 2009, 374, 1371–1382, doi:10.1016/s0140-6736(09)61338-6.

3. Webb, P.M.; Jordan, S.J. Epidemiology of epithelial ovarian cancer. Best practice & research. Clinical obstetrics & gynaecology 2017, 41, 3–14, doi:10.1016/j.bpobgyn.2016.08.006.

4. Gottesman, M.M. Mechanisms of cancer drug resistance. Annual review of medicine 2002, 53, 615–627, doi:10.1146/annurev.med.53.082901.103929.

5. Greenlee, R.T.; Murray, T.; Bolden, S.; Wingo, P.A. Cancer statistics, 2000. CA: a cancer journal for clinicians 2000, 50, 7–33, doi:10.3322/canjclin.50.1.7.

6. Gottesman, M.M.; Fojo, T.; Bates, S.E. Multidrug resistance in cancer: role of ATP-dependent transporters. Nature reviews. Cancer 2002, 2, 48–58, doi:10.1038/nrc706.

7. Leonard, G.D.; Fojo, T.; Bates, S.E. The role of ABC transporters in clinical practice. The oncologist 2003, 8, 411–424, doi:10.1634/theoncologist.8-5-411.

8. Mansoori, B.; Mohammadi, A.; Davudian, S.; Shirjang, S.; Baradaran, B. The Different Mechanisms of Cancer Drug Resistance: A Brief Review. Advanced pharmaceutical bulletin 2017, 7, 339–348, doi:10.15171/apb.2017.041.

9. Correia, A.L.; Bissell, M.J. The tumor microenvironment is a dominant force in multidrug resistance. Drug resistance updates: reviews and commentaries in antimicrobial and anticancer chemotherapy 2012, 15, 39–49, doi:10.1016/j.drup.2012.01.006.

10. Lee, R.C.; Feinbaum, R.L.; Ambros, V. The C. elegans heterochronic gene lin-4 encodes small RNAs with antisense complementarity to lin-14. Cell 1993, 75, 843–854, doi:10.1016/0092-8674(93)90529-y.

11. Bartel, D.P. MicroRNAs: genomics, biogenesis, mechanism, and function. Cell 2004, 116, 281–297, doi:10.1016/s0092-8674(04)00045-5.

12. Mihanfar, A.; Fattahi, A.; Nejabati, H.R. MicroRNA-mediated drug resistance in ovarian cancer. Journal of cellular physiology 2019, 234, 3180–3191, doi:10.1002/jcp.26060.

13. Baranwal, S.; Alahari, S.K. miRNA control of tumor cell invasion and metastasis. International journal of cancer 2010, 126, 1283–1290, doi:10.1002/ijc.25014.

14. Lee, Y.; Kim, M.; Han, J.; Yeom, K.H.; Lee, S.; Baek, S.H.; Kim, V.N. MicroRNA genes are transcribed by RNA polymerase II. The EMBO journal 2004, 23, 4051–4060, doi:10.1038/sj.emboj.7600385.

15. Kamanu, T.K.; Radovanovic, A.; Archer, J.A.; Bajic, V.B. Exploration of miRNA families for hypotheses generation. Scientific reports 2013, 3, 2940, doi:10.1038/srep02940.

16. Zou, Q.; Mao, Y.; Hu, L.; Wu, Y.; Ji, Z. miRClassify: an advanced web server for miRNA family classification and annotation. Computers in biology and medicine 2014, 45, 157–160, doi:10.1016/j.compbiomed.2013.12.007.

17. Zhang, W.; Dahlberg, J.E.; Tam, W. MicroRNAs in tumorigenesis: a primer. The American journal of pathology 2007, 171, 728–738, doi:10.2353/ajpath.2007.070070.

18. Lu, J.; Getz, G.; Miska, E.A.; Alvarez-Saavedra, E.; Lamb, J.; Peck, D.; Sweet-Cordero, A.; Ebert, B.L.; Mak, R.H.; Ferrando, A.A., et al. MicroRNA expression profiles classify human cancers. Nature 2005, 435, 834–838, doi:10.1038/nature03702.

19. Shenouda, S.K.; Alahari, S.K. MicroRNA function in cancer: oncogene or a tumor suppressor? Cancer metastasis reviews 2009, 28, 369–378, doi:10.1007/s10555-009-9188-5.

20. Lin, J.; Zhang, L.; Huang, H.; Huang, Y.; Huang, L.; Wang, J.; Huang, S.; He, L.; Zhou, Y.; Jia, W., et al. MiR-26b/KPNA2 axis inhibits epithelial ovarian carcinoma proliferation and metastasis through downregulating OCT4. Oncotarget 2015, 6, 23793–23806, doi:10.18632/oncotarget.4363.

21. Teng, Y.; Zhang, Y.; Qu, K.; Yang, X.; Fu, J.; Chen, W.; Li, X. MicroRNA-29B (mir-29b) regulates the Warburg effect in ovarian cancer by targeting AKT2 and AKT3. Oncotarget 2015, 6, 40799–40814, doi:10.18632/oncotarget.5695.

22. Sestito, R.; Cianfrocca, R.; Rosano, L.; Tocci, P.; Semprucci, E.; Di Castro, V.; Caprara, V.; Ferrandina, G.; Sacconi, A.; Blandino, G., et al. miR-30a inhibits endothelin A receptor and chemoresistance in ovarian carcinoma. Oncotarget 2016, 7, 4009–4023, doi:10.18632/oncotarget.6546.

23. Chen, X.; Chen, S.; Xiu, Y.L.; Sun, K.X.; Zong, Z.H.; Zhao, Y. RhoC is a major target of microRNA-93-5P in epithelial ovarian carcinoma tumorigenesis and progression. Molecular cancer 2015, 14, 31, doi:10.1186/s12943-015-0304-6.

24. Chen, S.; Chen, X.; Xiu, Y.L.; Sun, K.X.; Zhao, Y. Inhibition of Ovarian Epithelial Carcinoma Tumorigenesis and Progression by microRNA 106b Mediated through the RhoC Pathway. PloS one 2015, 10, e0125714, doi:10.1371/journal.pone.0125714.

25. Ying, X.; Wei, K.; Lin, Z.; Cui, Y.; Ding, J.; Chen, Y.; Xu, B. MicroRNA-125b Suppresses Ovarian Cancer Progression via Suppression of the Epithelial-Mesenchymal Transition Pathway by Targeting the SET Protein. Cellular physiology and biochemistry: international journal of experimental cellular physiology, biochemistry, and pharmacology 2016, 39, 501–510, doi:10.1159/000445642.

26. Chen, H.; Zhang, L.; Zhang, L.; Du, J.; Wang, H.; Wang, B. MicroRNA-183 correlates cancer prognosis, regulates cancer proliferation and bufalin sensitivity in epithelial ovarian caner. American journal of translational research 2016, 8, 1748–1755.

27. Yang, L.; Wei, Q.M.; Zhang, X.W.; Sheng, Q.; Yan, X.T. MiR-376a promotion of proliferation and metastases in ovarian cancer: Potential role as a biomarker. Life sciences 2017, 173, 62–67, doi:10.1016/j.lfs.2016.12.007.

28. Liu, J.; Dou, Y.; Sheng, M. Inhibition of microRNA-383 has tumor suppressive effect in human epithelial ovarian cancer through the action on caspase-2 gene. Biomedicine & pharmacotherapy = Biomedecine & pharmacotherapie 2016, 83, 1286–1294, doi:10.1016/j.biopha.2016.07.038.

29. Chaluvally-Raghavan, P.; Jeong, K.J.; Pradeep, S.; Silva, A.M.; Yu, S.; Liu, W.; Moss, T.; Rodriguez-Aguayo, C.; Zhang, D.; Ram, P., et al. Direct Upregulation of STAT3 by MicroRNA-551b-3p Deregulates Growth and Metastasis of Ovarian Cancer. Cell reports 2016, 15, 1493–1504, doi:10.1016/j.celrep.2016.04.034.

30. Zhang, X.; Liu, J.; Zang, D.; Wu, S.; Liu, A.; Zhu, J.; Wu, G.; Li, J.; Jiang, L. Upregulation of miR-572 transcriptionally suppresses SOCS1 and p21 and contributes to human ovarian cancer progression. Oncotarget 2015, 6, 15180–15193, doi:10.18632/oncotarget.3737.

31. Su, L.; Liu, M. Correlation analysis on the expression levels of microRNA-23a and microRNA-23b and the incidence and prognosis of ovarian cancer. Oncology letters 2018, 16, 262–266, doi:10.3892/ol.2018.8669.

32. Kim, Y.W.; Kim, E.Y.; Jeon, D.; Liu, J.L.; Kim, H.S.; Choi, J.W.; Ahn, W.S. Differential microRNA expression signatures and cell type-specific association with Taxol resistance in ovarian cancer cells. Drug design, development and therapy 2014, 8, 293–314, doi:10.2147/dddt.s51969.

33. Mitamura, T.; Watari, H.; Wang, L.; Kanno, H.; Hassan, M.K.; Miyazaki, M.; Katoh, Y.; Kimura, T.; Tanino, M.; Nishihara, H., et al. Downregulation of miRNA-31 induces taxane resistance in ovarian cancer cells through increase of receptor tyrosine kinase MET. Oncogenesis 2013, 2, e40, doi:10.1038/oncsis.2013.3.

34. Wang, Y.; Bao, W.; Liu, Y.; Wang, S.; Xu, S.; Li, X.; Li, Y.; Wu, S. miR-98-5p contributes to cisplatin resistance in epithelial ovarian cancer by suppressing miR-152 biogenesis via targeting Dicer1. Cell death & disease 2018, 9, 447, doi:10.1038/s41419-018-0390-7.

35. Wang, H.; Ren, S.; Xu, Y.; Miao, W.; Huang, X.; Qu, Z.; Li, J.; Liu, X.; Kong, P. MicroRNA-195 reverses the resistance to temozolomide through targeting cyclin E1 in glioma cells. Anti-cancer drugs 2019, 30, 81–88, doi:10.1097/cad.0000000000000700.

36. Chen, L.Z.; Ding, Z.; Zhang, Y.; He, S.T.; Wang, X.H. MiR-203 over-expression promotes prostate cancer cell apoptosis and reduces ADM resistance. European review for medical and pharmacological sciences 2018, 22, 3734–3741, doi:10.26355/eurrev_201806_15253.

37. Januchowski, R.; Wojtowicz, K.; Sujka-Kordowska, P.; Andrzejewska, M.; Zabel, M. MDR gene expression analysis of six drug-resistant ovarian cancer cell lines. BioMed research international 2013, 2013, 241763, doi:10.1155/2013/241763.

38. Stelcer, E.; Kulcenty, K.; Rucinski, M.; Jopek, K.; Richter, M.; Trzeciak, T.; Suchorska, W.M. The Role of MicroRNAs in Early Chondrogenesis of Human Induced Pluripotent Stem Cells (hiPSCs). International journal of molecular sciences 2019, 20, doi:10.3390/ijms20184371.

39. Kulcenty, K.; Wroblewska, J.P.; Rucinski, M.; Kozlowska, E.; Jopek, K.; Suchorska, W.M. MicroRNA Profiling During Neural Differentiation of Induced Pluripotent Stem Cells. International journal of molecular sciences 2019, 20, doi:10.3390/ijms20153651.

40. Gautier, L.; Cope, L.; Bolstad, B.M.; Irizarry, R.A. affy--analysis of Affymetrix GeneChip data at the probe level. Bioinformatics (Oxford, England) 2004, 20, 307–315, doi:10.1093/bioinformatics/btg405.

41. Ritchie, M.E.; Phipson, B.; Wu, D.; Hu, Y.; Law, C.W.; Shi, W.; Smyth, G.K. limma powers differential expression analyses for RNA-sequencing and microarray studies. Nucleic acids research 2015, 43, e47, doi:10.1093/nar/gkv007.

42. Cava, C.; Colaprico, A.; Bertoli, G.; Graudenzi, A.; Silva, T.C.; Olsen, C.; Noushmehr, H.; Bontempi, G.; Mauri, G.; Castiglioni, I. SpidermiR: An R/Bioconductor Package for Integrative Analysis with miRNA Data. International journal of molecular sciences 2017, 18, doi:10.3390/ijms18020274.

43. Januchowski, R.; Sterzynska, K.; Zawierucha, P.; Rucinski, M.; Swierczewska, M.; Partyka, M.; Bednarek-Rajewska, K.; Brazert, M.; Nowicki, M.; Zabel, M., et al. Microarray-based detection and expression analysis of new genes associated with drug resistance in ovarian cancer cell lines. Oncotarget 2017, 8, 49944–49958, doi:10.18632/oncotarget.18278.

44. Januchowski, R.; Zawierucha, P.; Rucinski, M.; Zabel, M. Microarray-based detection and expression analysis of extracellular matrix proteins in drugresistant ovarian cancer cell lines. Oncology reports 2014, 32, 1981–1990, doi:10.3892/or.2014.3468.

45. Januchowski, R.; Zawierucha, P.; Andrzejewska, M.; Rucinski, M.; Zabel, M. Microarray-based detection and expression analysis of ABC and SLC transporters in drug-resistant ovarian cancer cell lines. Biomedicine & pharmacotherapy = Biomedecine & pharmacotherapie 2013, 67, 240–245, doi:10.1016/j.biopha.2012.11.011.

46. Shannon, P.; Markiel, A.; Ozier, O.; Baliga, N.S.; Wang, J.T.; Ramage, D.; Amin, N.; Schwikowski, B.; Ideker, T. Cytoscape: a software environment for integrated models of biomolecular interaction networks. Genome research 2003, 13, 2498–2504, doi:10.1101/gr.1239303.

47. Ma, J.; Dong, C.; Ji, C. MicroRNA and drug resistance. Cancer gene therapy 2010, 17, 523–531, doi:10.1038/cgt.2010.18.

48. Hassan, M.K.; Watari, H.; Mitamura, T.; Mohamed, Z.; El-Khamisy, S.F.; Ohba, Y.; Sakuragi, N. P18/Stathmin1 is regulated by miR-31 in ovarian cancer in response to taxane. Oncoscience 2015, 2, 294–308, doi:10.18632/oncoscience.143.

49. Li, M.; Chen, W.; Zhang, H.; Zhang, Y.; Ke, F.; Wu, X.; Zhang, Y.; Weng, M.; Liu, Y.; Gong, W. MiR-31 regulates the cisplatin resistance by targeting Src in gallbladder cancer. Oncotarget 2016, 7, 83060–83070, doi:10.18632/oncotarget.13067.

50. Januchowski, R.; Wojtowicz, K.; Sterzyska, K.; Sosiska, P.; Andrzejewska, M.; Zawierucha, P.; Nowicki, M.; Zabel, M. Inhibition of ALDH1A1 activity decreases expression of drug transporters and reduces chemotherapy resistance in ovarian cancer cell lines. The international journal of biochemistry & cell biology 2016, 78, 248–259, doi:10.1016/j.biocel.2016.07.017.

51. Sterzynska, K.; Klejewski, A.; Wojtowicz, K.; Swierczewska, M.; Nowacka, M.; Kazmierczak, D.; Andrzejewska, M.; Rusek, D.; Brazert, M.; Brazert, J., et al. Mutual Expression of ALDH1A1, LOX, and Collagens in Ovarian Cancer Cell Lines as Combined CSCs- and ECM-Related Models of Drug Resistance Development. International journal of molecular sciences 2018, 20, doi:10.3390/ijms20010054.

52. Lv, C.; Li, F.; Li, X.; Tian, Y.; Zhang, Y.; Sheng, X.; Song, Y.; Meng, Q.; Yuan, S.; Luan, L., et al. MiR-31 promotes mammary stem cell expansion and breast tumorigenesis by suppressing Wnt signaling antagonists. Nature communications 2017, 8, 1036, doi:10.1038/s41467-017-01059-5.

53. Yang, M.; Shan, X.; Zhou, X.; Qiu, T.; Zhu, W.; Ding, Y.; Shu, Y.; Liu, P. miR-1271 regulates cisplatin resistance of human gastric cancer cell lines by targeting IGF1R, IRS1, mTOR, and BCL2. Anti-cancer agents in medicinal chemistry 2014, 14, 884–891, doi:10.2174/1871520614666140528161318.

54. Yang, L.; Wang, Y.; Li, Y.J.; Zeng, C.C. Chemo-resistance of A172 glioblastoma cells is controlled by miR-1271-regulated Bcl-2. Biomedicine & pharmacotherapy = Biomedecine & pharmacotherapie 2018, 108, 734–740, doi:10.1016/j.biopha.2018.08.102.

55. Lu, G.; Fu, D.; Jia, C.; Chai, L.; Han, Y.; Liu, J.; Wu, T.; Xie, R.; Chang, Z.; Yang, H., et al. Reduced miR-105-1 levels are associated with poor survival of patients with non-small cell lung cancer. Oncology letters 2017, 14, 7842–7848, doi:10.3892/ol.2017.7228.

56. Li, H.Y.; Liang, J.L.; Kuo, Y.L.; Lee, H.H.; Calkins, M.J.; Chang, H.T.; Lin, F.C.; Chen, Y.C.; Hsu, T.I.; Hsiao, M., et al. miR-105/93-3p promotes chemoresistance and circulating miR-105/93-3p acts as a diagnostic biomarker for triple negative breast cancer. Breast cancer research: BCR 2017, 19, 133, doi:10.1186/s13058-017-0918-2.

57. Xiang, Y.; Ma, N.; Wang, D.; Zhang, Y.; Zhou, J.; Wu, G.; Zhao, R.; Huang, H.; Wang, X.; Qiao, Y., et al. MiR-152 and miR-185 co-contribute to ovarian cancer cells cisplatin sensitivity by targeting DNMT1 directly: a novel epigenetic therapy independent of decitabine. Oncogene 2014, 33, 378–386, doi:10.1038/onc.2012.575.

58. Dai, J.; Wei, R.; Zhang, P.; Kong, B. Overexpression of microRNA-195-5p reduces cisplatin resistance and angiogenesis in ovarian cancer by inhibiting the PSAT1-dependent GSK3beta/beta-catenin signaling pathway. Journal of translational medicine 2019, 17, 190, doi:10.1186/s12967-019-1932-1.

59. Xu, C.Z.; Xie, J.; Jin, B.; Chen, X.W.; Sun, Z.F.; Wang, B.X.; Dong, P. Gene and microRNA expression reveals sensitivity to paclitaxel in laryngeal cancer cell line. International journal of clinical and experimental pathology 2013, 6, 1351–1361.

60. Dai, Y.; Xie, C.H.; Neis, J.P.; Fan, C.Y.; Vural, E.; Spring, P.M. MicroRNA expression profiles of head and neck squamous cell carcinoma with docetaxel-induced multidrug resistance. Head & neck 2011, 33, 786–791, doi:10.1002/hed.21540.

61. Sun, W.; Ma, Y.; Chen, P.; Wang, D. MicroRNA-10a silencing reverses cisplatin resistance in the A549/cisplatin human lung cancer cell line via the transforming growth factor-beta/Smad2/STAT3/STAT5 pathway. Molecular medicine reports 2015, 11, 3854–3859, doi:10.3892/mmr.2015.3181.

62. Yan, M.; Yang, X.; Shen, R.; Wu, C.; Wang, H.; Ye, Q.; Yang, P.; Zhang, L.; Chen, M.; Wan, B., et al. miR-146b promotes cell proliferation and increases chemosensitivity, but attenuates cell migration and invasion via FBXL10 in ovarian cancer. Cell death & disease 2018, 9, 1123, doi:10.1038/s41419-018-1093-9.

63. Cheng, R.; Lu, C.; Zhang, G.; Zhang, G.; Zhao, G. Overexpression of miR-203 increases the sensitivity of NSCLC A549/H460 cell lines to cisplatin by targeting Dickkopf-1. Oncology reports 2017, 37, 2129–2136, doi:10.3892/or.2017.5505.

64. Liu, Y.; Gao, S.; Chen, X.; Liu, M.; Mao, C.; Fang, X. Overexpression of miR-203 sensitizes paclitaxel (Taxol)-resistant colorectal cancer cells through targeting the salt-inducible kinase 2 (SIK2). Tumour biology: the journal of the International Society for Oncodevelopmental Biology and Medicine 2016, 37, 12231–12239, doi:10.1007/s13277-016-5066-2.

65. Mohamed, Z.; Hassan, M.K.; Okasha, S.; Mitamura, T.; Keshk, S.; Konno, Y.; Kato, T.; El-Khamisy, S.F.; Ohba, Y.; Watari, H. miR-363 confers taxane resistance in ovarian cancer by targeting the Hippo pathway member, LATS2. Oncotarget 2018, 9, 30053–30065, doi:10.18632/oncotarget.25698.

66. Ou, Y.; Zhai, D.; Wu, N.; Li, X. Downregulation of miR-363 increases drug resistance in cisplatin-treated HepG2 by dysregulating Mcl-1. Gene 2015, 572, 116–122, doi:10.1016/j.gene.2015.07.002.

67. Cao, L.; Wan, Q.; Li, F.; Tang, C.E. MiR-363 inhibits cisplatin chemoresistance of epithelial ovarian cancer by regulating snail-induced epithelial-mesenchymal transition. BMB reports 2018, 51, 456–461.

68. Yu, P.N.; Yan, M.D.; Lai, H.C.; Huang, R.L.; Chou, Y.C.; Lin, W.C.; Yeh, L.T.; Lin, Y.W. Downregulation of miR-29 contributes to cisplatin resistance of ovarian cancer cells. International journal of cancer 2014, 134, 542–551, doi:10.1002/ijc.28399.

69. Yang, L.; Li, N.; Yan, Z.; Li, C.; Zhao, Z. MiR-29a-Mediated CD133 Expression Contributes to Cisplatin Resistance in CD133(+) Glioblastoma Stem Cells. Journal of molecular neuroscience: MN 2018, 66, 369–377, doi:10.1007/s12031-018-1177-0.

70. Xu, W.; Li, Z.; Zhu, X.; Xu, R.; Xu, Y. miR-29 Family Inhibits Resistance to Methotrexate and Promotes Cell Apoptosis by Targeting COL3A1 and MCL1 in Osteosarcoma. Medical science monitor: international medical journal of experimental and clinical research 2018, 24, 8812–8821, doi:10.12659/msm.911972.

71. Sorrentino, A.; Liu, C.G.; Addario, A.; Peschle, C.; Scambia, G.; Ferlini, C. Role of microRNAs in drug-resistant ovarian cancer cells. Gynecologic oncology 2008, 111, 478–486, doi:10.1016/j.ygyno.2008.08.017.

72. Guo, Y.; Zhang, H.; Xie, D.; Hu, X.; Song, R.; Zhu, L. Non-coding RNA NEAT1/miR-214-3p contribute to doxorubicin resistance of urothelial bladder cancer preliminary through the Wnt/beta-catenin pathway. Cancer management and research 2018, 10, 4371–4380, doi:10.2147/cmar.s171126.

73. Zhang, Y.; Xu, W.; Ni, P.; Li, A.; Zhou, J.; Xu, S. MiR-99a and MiR-491 Regulate Cisplatin Resistance in Human Gastric Cancer Cells by Targeting CAPNS1. International journal of biological sciences 2016, 12, 1437–1447, doi:10.7150/ijbs.16529.

74. Akbari Moqadam, F.; Lange-Turenhout, E.A.; Aries, I.M.; Pieters, R.; den Boer, M.L. MiR-125b, miR-100 and miR-99a co-regulate vincristine resistance in childhood acute lymphoblastic leukemia. Leukemia research 2013, 37, 1315–1321, doi:10.1016/j.leukres.2013.06.027.

75. Zhu, X.; Li, Y.; Xie, C.; Yin, X.; Liu, Y.; Cao, Y.; Fang, Y.; Lin, X.; Xu, Y.; Xu, W., et al. miR-145 sensitizes ovarian cancer cells to paclitaxel by targeting Sp1 and Cdk6. International journal of cancer 2014, 135, 1286–1296, doi:10.1002/ijc.28774.

76. Khordadmehr, M.; Shahbazi, R.; Sadreddini, S.; Baradaran, B. miR-193: A new weapon against cancer. Journal of cellular physiology 2019, 234, 16861–16872, doi:10.1002/jcp.28368.

77. Janikova, M.; Zizkova, V.; Skarda, J.; Kharaishvili, G.; Radova, L.; Kolar, Z. Prognostic significance of miR-23b in combination with P-gp, MRP and LRP/MVP expression in non-small cell lung cancer. Neoplasma 2016, 63, 576–587, doi:10.4149/neo_2016_411.

78. Xu, Y.; Han, Y.F.; Ye, B.; Zhang, Y.L.; Dong, J.D.; Zhu, S.J.; Chen, J. miR-27b-3p is Involved in Doxorubicin Resistance of Human Anaplastic Thyroid Cancer Cells via Targeting Peroxisome Proliferator-Activated Receptor Gamma. Basic & clinical pharmacology & toxicology 2018, 123, 670–677, doi:10.1111/bcpt.13076.

79. Husted, S.; Sokilde, R.; Rask, L.; Cirera, S.; Busk, P.K.; Eriksen, J.; Litman, T. MicroRNA expression profiles associated with development of drug resistance in Ehrlich ascites tumor cells. Molecular pharmaceutics 2011, 8, 2055–2062, doi:10.1021/mp200255d.

80. Xu, X.; Jiang, X.; Chen, L.; Zhao, Y.; Huang, Z.; Zhou, H.; Shi, M. MiR-181a Promotes Apoptosis and Reduces Cisplatin Resistance by Inhibiting Osteopontin in Cervical Cancer Cells. Cancer biotherapy & radiopharmaceuticals 2019, 34, 559–565, doi:10.1089/cbr.2019.2858.

81. Wang, X.; Chen, X.; Meng, Q.; Jing, H.; Lu, H.; Yang, Y.; Cai, L.; Zhao, Y. MiR-181b regulates cisplatin chemosensitivity and metastasis by targeting TGFbetaR1/Smad signaling pathway in NSCLC. Scientific reports 2015, 5, 17618, doi:10.1038/srep17618.

82. Cheng, W.; Liu, T.; Wan, X.; Gao, Y.; Wang, H. MicroRNA-199a targets CD44 to suppress the tumorigenicity and multidrug resistance of ovarian cancer-initiating cells. The FEBS journal 2012, 279, 2047–2059, doi:10.1111/j.1742-4658.2012.08589.x.

83. Liu, M.X.; Siu, M.K.; Liu, S.S.; Yam, J.W.; Ngan, H.Y.; Chan, D.W. Epigenetic silencing of microRNA-199b-5p is associated with acquired chemoresistance via activation of JAG1-Notch1 signaling in ovarian cancer. Oncotarget 2014, 5, 944–958, doi:10.18632/oncotarget.1458.

84. Januchowski, R.; Zawierucha, P.; Rucinski, M.; Andrzejewska, M.; Wojtowicz, K.; Nowicki, M.; Zabel, M. Drug transporter expression profiling in chemoresistant variants of the A2780 ovarian cancer cell line. Biomedicine & pharmacotherapy = Biomedecine & pharmacotherapie 2014, 68, 447–453, doi:10.1016/j.biopha.2014.02.002.

85. Januchowski, R.; Zawierucha, P.; Rucinski, M.; Nowicki, M.; Zabel, M. Extracellular matrix proteins expression profiling in chemoresistant variants of the A2780 ovarian cancer cell line. BioMed research international 2014, 2014, 365867, doi:10.1155/2014/365867.

86. Prislei, S.; Mozzetti, S.; Filippetti, F.; De Donato, M.; Raspaglio, G.; Cicchillitti, L.; Scambia, G.; Ferlini, C. From plasma membrane to cytoskeleton: a novel function for semaphorin 6A. Molecular cancer therapeutics 2008, 7, 233–241, doi:10.1158/1535-7163.mct-07-0390.

87. Huang, H.; Chen, J.; Ding, C.M.; Jin, X.; Jia, Z.M.; Peng, J. LncRNA NR2F1-AS1 regulates hepatocellular carcinoma oxaliplatin resistance by targeting ABCC1 via miR-363. Journal of cellular and molecular medicine 2018, 22, 3238–3245, doi:10.1111/jcmm.13605.

88. Januchowski, R.; Swierczewska, M.; Sterzynska, K.; Wojtowicz, K.; Nowicki, M.; Zabel, M. Increased Expression of Several Collagen Genes is Associated with Drug Resistance in Ovarian Cancer Cell Lines. Journal of Cancer 2016, 7, 1295–1310, doi:10.7150/jca.15371.

89. Swierczewska, M.; Sterzynska, K.; Wojtowicz, K.; Kazmierczak, D.; Izycki, D.; Nowicki, M.; Zabel, M.; Januchowski, R. PTPRK Expression Is Downregulated in Drug Resistant Ovarian Cancer Cell Lines, and Especially in ALDH1A1 Positive CSCs-Like Populations. International journal of molecular sciences 2019, 20, doi:10.3390/ijms20082053.

90. Zou, J.; Yin, F.; Wang, Q.; Zhang, W.; Li, L. Analysis of microarray-identified genes and microRNAs associated with drug resistance in ovarian cancer. International journal of clinical and experimental pathology 2015, 8, 6847–6858.

91. Zhou, R. The Eph family receptors and ligands. Pharmacology & therapeutics 1998, 77, 151–181, doi:10.1016/s0163-7258(97)00112-5.

92. Zhang, S.J.; Zhang, G.; Zhao, Y.F.; Wu, Y.; Li, J.; Chai, Y.X. [Expression of EphA7 protein in primary hepatocellular carcinoma and its clinical significance]. Zhonghua wai ke za zhi [Chinese journal of surgery] 2010, 48, 53–56.

93. Giani, F.; Russo, G.; Pennisi, M.; Sciacca, L.; Frasca, F.; Pappalardo, F. Computational modeling reveals MAP3K8 as mediator of resistance to vemurafenib in thyroid cancer stem cells. Bioinformatics (Oxford, England) 2019, 35, 2267–2275, doi:10.1093/bioinformatics/bty969.

94. Izycka, N.; Sterzynska, K.; Januchowski, R.; Nowak-Markwitz, E. Semaphorin 3A (SEMA3A), protocadherin 9 (PCdh9), and S100 calcium binding protein A3 (S100A3) as potential biomarkers of carcinogenesis and chemoresistance of different neoplasms, including ovarian cancer - review of literature. Ginekologia polska 2019, 90, 223–227, doi:10.5603/gp.2019.0040.

95. Tang, C.; Gao, X.; Liu, H.; Jiang, T.; Zhai, X. Decreased expression of SEMA3A is associated with poor prognosis in gastric carcinoma. International journal of clinical and experimental pathology 2014, 7, 4782–4794.

96. Jiang, H.; Qi, L.; Wang, F.; Sun, Z.; Huang, Z.; Xi, Q. Decreased semaphorin 3A expression is associated with a poor prognosis in patients with epithelial ovarian carcinoma. International journal of molecular medicine 2015, 35, 1374–1380, doi:10.3892/ijmm.2015.2142.

97. Swierczewska, M.; Klejewski, A.; Brazert, M.; Kazmierczak, D.; Izycki, D.; Nowicki, M.; Zabel, M.; Januchowski, R. New and Old Genes Associated with Primary and Established Responses to Paclitaxel Treatment in Ovarian Cancer Cell Lines. Molecules (Basel, Switzerland) 2018, 23, doi:10.3390/molecules23040891.

98. Jin, Y.; Hong, F.; Bao, Q.; Xu, Q.; Duan, R.; Zhu, Z.; Zhang, W.; Ma, C. MicroRNA-145 suppresses osteogenic differentiation of human jaw bone marrow mesenchymal stem cells partially via targeting semaphorin 3A. Connective tissue research 2019, 10.1080/03008207.2019.1643334, 1–9, doi:10.1080/03008207.2019.1643334.

99. Wang, F.; Xia, J.; Wang, N.; Zong, H. miR-145 inhibits proliferation and invasion of esophageal squamous cell carcinoma in part by targeting c-Myc. Onkologie 2013, 36, 754–758, doi:10.1159/000356978.

100. Li, J.; Li, X.; Wu, L.; Pei, M.; Li, H.; Jiang, Y. miR-145 inhibits glutamine metabolism through c-myc/GLS1 pathways in ovarian cancer cells. Cell biology international 2019, 43, 921–930, doi:10.1002/cbin.11182.

101. Nadiminty, N.; Tummala, R.; Lou, W.; Zhu, Y.; Zhang, J.; Chen, X.; eVere White, R.W.; Kung, H.J.; Evans, C.P.; Gao, A.C. MicroRNA let-7c suppresses androgen receptor expression and activity via regulation of Myc expression in prostate cancer cells. The Journal of biological chemistry 2012, 287, 1527–1537, doi:10.1074/jbc.M111.278705.

102. Frank, M.; Kemler, R. Protocadherins. Current opinion in cell biology 2002, 14, 557–562, doi:10.1016/s0955-0674(02)00365-4.

103. Kim, S.Y.; Yasuda, S.; Tanaka, H.; Yamagata, K.; Kim, H. Non-clustered protocadherin. Cell adhesion & migration 2011, 5, 97–105, doi:10.4161/cam.5.2.14374.

104. Shi, C.; Zhang, Z. Screening of potentially crucial genes and regulatory factors involved in epithelial ovarian cancer using microarray analysis. Oncology letters 2017, 14, 725–732, doi:10.3892/ol.2017.6183.

105. Chen, Y.; Xiang, H.; Zhang, Y.; Wang, J.; Yu, G. Loss of PCDH9 is associated with the differentiation of tumor cells and metastasis and predicts poor survival in gastric cancer. Clinical & experimental metastasis 2015, 32, 417–428, doi:10.1007/s10585-015-9712-7.

106. Sofeu Feugaing, D.D.; Gotte, M.; Viola, M. More than matrix: the multifaceted role of decorin in cancer. European journal of cell biology 2013, 92, 1–11, doi:10.1016/j.ejcb.2012.08.004.

107. Kwan, P.; Ding, J.; Tredget, E.E. MicroRNA 181b regulates decorin production by dermal fibroblasts and may be a potential therapy for hypertrophic scar. PloS one 2015, 10, e0123054, doi:10.1371/journal.pone.0123054.

108. Lucero, H.A.; Kagan, H.M. Lysyl oxidase: an oxidative enzyme and effector of cell function. Cellular and molecular life sciences: CMLS 2006, 63, 2304–2316, doi:10.1007/s00018-006-6149-9.

109. Erler, J.T.; Bennewith, K.L.; Nicolau, M.; Dornhofer, N.; Kong, C.; Le, Q.T.; Chi, J.T.; Jeffrey, S.S.; Giaccia, A.J. Lysyl oxidase is essential for hypoxia-induced metastasis. Nature 2006, 440, 1222–1226, doi:10.1038/nature04695.

110. Ji, F.; Wang, Y.; Qiu, L.; Li, S.; Zhu, J.; Liang, Z.; Wan, Y.; Di, W. Hypoxia inducible factor 1alpha-mediated LOX expression correlates with migration and invasion in epithelial ovarian cancer. International journal of oncology 2013, 42, 1578–1588, doi:10.3892/ijo.2013.1878.

111. Ween, M.P.; Oehler, M.K.; Ricciardelli, C. Transforming growth Factor-Beta-Induced Protein (TGFBI)/(betaig-H3): a matrix protein with dual functions in ovarian cancer. International journal of molecular sciences 2012, 13, 10461–10477, doi:10.3390/ijms130810461.

112. Karlan, B.Y.; Dering, J.; Walsh, C.; Orsulic, S.; Lester, J.; Anderson, L.A.; Ginther, C.L.; Fejzo, M.; Slamon, D. POSTN/TGFBI-associated stromal signature predicts poor prognosis in serous epithelial ovarian cancer. Gynecologic oncology 2014, 132, 334–342, doi:10.1016/j.ygyno.2013.12.021.

113. Bhushan, R.; Grunhagen, J.; Becker, J.; Robinson, P.N.; Ott, C.E.; Knaus, P. miR-181a promotes osteoblastic differentiation through repression of TGF-beta signalling molecules. The international journal of biochemistry & cell biology 2013, 45, 696–705, doi:10.1016/j.biocel.2012.12.008.

114. Young, M.F.; Kerr, J.M.; Termine, J.D.; Wewer, U.M.; Wang, M.G.; McBride, O.W.; Fisher, L.W. cDNA cloning, mRNA distribution and heterogeneity, chromosomal location, and RFLP analysis of human osteopontin (OPN). Genomics 1990, 7, 491–502, doi:10.1016/0888-7543(90)90191-v.

115. Kim, J.H.; Skates, S.J.; Uede, T.; Wong, K.K.; Schorge, J.O.; Feltmate, C.M.; Berkowitz, R.S.; Cramer, D.W.; Mok, S.C. Osteopontin as a potential diagnostic biomarker for ovarian cancer. Jama 2002, 287, 1671–1679, doi:10.1001/jama.287.13.1671.

116. Bao, L.H.; Sakaguchi, H.; Fujimoto, J.; Tamaya, T. Osteopontin in metastatic lesions as a prognostic marker in ovarian cancers. Journal of biomedical science 2007, 14, 373–381, doi:10.1007/s11373-006-9143-1.

117. Das, S.; Samant, R.S.; Shevde, L.A. Nonclassical activation of Hedgehog signaling enhances multidrug resistance and makes cancer cells refractory to Smoothened-targeting Hedgehog inhibition. The Journal of biological chemistry 2013, 288, 11824–11833, doi:10.1074/jbc.M112.432302.

118. Cui, R.; Takahashi, F.; Ohashi, R.; Yoshioka, M.; Gu, T.; Tajima, K.; Unnoura, T.; Iwakami, S.; Hirama, M.; Ishiwata, T., et al. Osteopontin is involved in the formation of malignant pleural effusion in lung cancer. Lung cancer (Amsterdam, Netherlands) 2009, 63, 368–374, doi:10.1016/j.lungcan.2008.06.020.

119. Huo, X.; Zhang, K.; Yi, L.; Mo, Y.; Liang, Y.; Zhao, J.; Zhang, Z.; Xu, Y.; Zhen, G. Decreased epithelial and plasma miR-181b-5p expression associates with airway eosinophilic inflammation in asthma. Clinical and experimental allergy: journal of the British Society for Allergy and Clinical Immunology 2016, 46, 1281–1290, doi:10.1111/cea.12754.

120. Helleman, J.; Jansen, M.P.; Span, P.N.; van Staveren, I.L.; Massuger, L.F.; Meijer-van Gelder, M.E.; Sweep, F.C.; Ewing, P.C.; van der Burg, M.E.; Stoter, G., et al. Molecular profiling of platinum resistant ovarian cancer. International journal of cancer 2006, 118, 1963–1971, doi:10.1002/ijc.21599.

